# MicroRNA-181 influences Alzheimer’s risk by regulating neprilysin and microtubule-associated tau pathways, offering a novel target

**DOI:** 10.64898/2026.06.11.731747

**Authors:** Ruizhi Wang, Bryan Maloney, Kwangsik Nho, John S. Beck, Scott E. Counts, Debomoy K. Lahiri

## Abstract

Alzheimer’s disease (AD) is characterized by amyloid-β (Aβ) peptide plaques and neurofibrillary tangles from hyperphosphorylated tau, though factors linking amyloid and tau pathology remain unclear. We investigated whether microRNA-181d-5p (miR-181d) associates with AD-related brain changes and regulates neprilysin and tau. Modeling miR-181d across individuals with no cognitive impairment, mild cognitive impairment, and AD revealed region- and sex-specific associations. Higher miR-181d levels associated with greater AD probability in the temporal lobe and cerebellum, and lower probability in the posterior cingulate cortex of males; miR-181c attenuated these probabilities. SNPs near MIR181 associated with altered entorhinal cortical thickness. In cellular models, miR-181 reduced neprilysin 3′-UTR activity, mRNA, protein, and enzymatic activity, while increasing tau mRNA and protein. Neprilysin diminution impairs Aβ clearance and elevates tau, contributing to AD. RNA sequencing identified miR-181d-responsive neurodegenerative pathways. These findings identify miR-181 as a regulator of AD-relevant amyloid and tau pathways, providing novel targets.

**Teaser:** MiRNA-181 is a key regulator of Alzheimer’s risk through its effects on neprilysin and tau proteins, a novel potential target.

## Introduction

Alzheimer’s disease (AD), the most common cause of dementia worldwide, is histopathologically characterized by amyloid plaques and neurofibrillary tangles in vulnerable brain regions. Amyloid plaques are formed by abnormal aggregation of extracellular amyloid β (Aβ) peptides, primarily Aβ40 and Aβ42. Neurofibrillary tangles are mainly composed of hyperphosphorylated tau proteins. However, mechanisms linking these pathological changes to disease state remain incompletely understood.

Aβ is derived from amyloid precursor protein (APP) via sequential cleavage by β- and γ-secretases. Several enzymes can degrade Aβ, including neprilysin (also known as membrane metallo-endopeptidase [MME]), CD10 or neutral endopeptidase (NEP) (*1*), angiotensin-converting enzyme (ACE) (*2*), endothelin-converting enzyme (ECE) (*3*), and insulin-degrading enzyme (IDE) (*4*). Neprilysin is capable of degrading intracellular and extracellular Aβ, most actively at neutral pH, and is relevant to AD-associated metabolism (*5*).

MicroRNAs (miRNAs) are a group of small non-coding RNAs. miRNAs typically regulate gene and protein expression by reducing target mRNA stability and translation. In the canonical pathway, miRNA as a component of RNA-induced silencing complex (RISC) mediates target recognition via base paring with the target mRNA 3’ untranslated region (UTR). After binding to target mRNA 3’ UTR, RISC can degrade target mRNA or inhibit target mRNA translation (*6–8*).

The miR-181 family comprises multiple mature miRNAs, including four “5p” members, specifically miR-181a-5p, miR-181b-5p, miR-181c-5p and miR-181d-5p. These are encoded by six genes, specifically *MIR181A1, MIR181A2, MIR181B1, MIR181B2, MIR181C,* and *MIR181D* derived from six different sequences on three chromosomes (1, 9 and 19). The mature miR-181 miRNA sequences are well conserved among these genes, sharing the same seed sequence and great similarity in target binding. The miR-181 family is involved in multiple biological and pathological processes (*9*). miR-181 is dysregulated in brains, cerebrospinal fluid (CSF) and blood of human AD patients (*10–12*). miR-181 is up-regulated in the hippocampus of 3xTg AD mice (*13*). miR-181c reduces expression of serine palmitoyltransferase (SPT), a rate-limiting enzyme in ceramide synthesis, which in turn reduces Aβ levels (*14*).

Herein, we quantified miR-181a-d in the temporal lobe, cerebellum and posterior cingulate cortex (PCC) of human brain specimens from postmortem no cognitive impairment (NCI), mild cognitive impairment (MCI), and AD patients. Notably, miR-181d-5p, but not miR-181 a, b or c, was significantly associated with AD diagnosis probability. In neuronal cell models, miR-181d-5p reduced neprilysin 3′-UTR reporter activity and reduced MME mRNA, neprilysin protein, and neprilysin enzyme activity.. Moreover, miR-181d transfection increased tau mRNA and protein expression as well as phosphorylation. A single nucleotide polymorphism (SNP) in close proximity to *MIR181A1* and *MIR181B1* genes was associated with the thickness of entorhinal cortex. RNA sequencing data revealed miR-181d-responsive genes were enriched in Alzheimer’s disease and other neurodegenerative disease pathways. In short, our results presented here suggest that miRNA-181 as a key regulator of AD risk through its cooperative effects on neprilysin and microtubule-associated tau proteins, influencing both amyloid and tau pathways and highlighting its potential as a novel target.

## Results

### Brain miR-181d levels associated with altered AD diagnosis probability

When we modeled levels of miR-181d vs NCI, MCI, and AD diagnosis in brain samples, we found a significant positive association between elevated miR-181d levels and increased diagnosis probability of AD in the temporal lobe and cerebellum (Fig. 1A-F). However, in the posterior cingulate cortex (PCC) of male patients, miR-181d is associated with reduced AD probability. We tested miR-181a, b, and c as covariates and found that of the three, miR-181c interacted with miR-181d and reduced the mir-181d associated probability of AD. We also found that the miR-181d effect was stronger in female than in male subjects. miR-181a, b, and c levels were very highly correlated with each other (Supplemental Fig. 1).

**Fig. 1.**
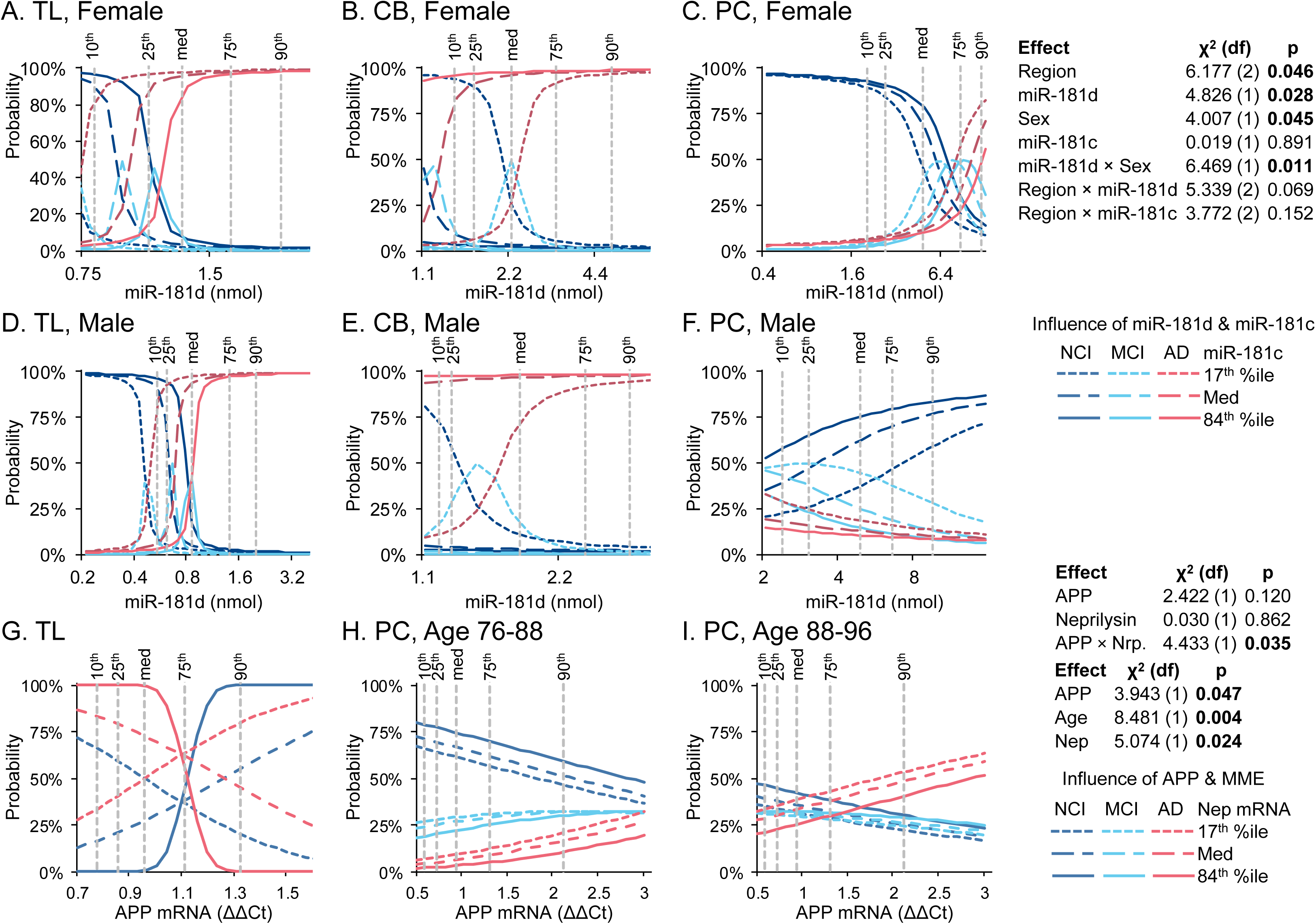
Effects of miR-181d/c and APP mRNA levels in brain samples on probability of AD. Total RNA was extracted from human brain samples and subject to qRT-PCR quantification (X-axis), as described in the proposal. We constructed a logistic regression model that also considered effects of age, sex and brain region. Models revealed that each variable exerted significant effect on probability of AD (Y-axis). These were used to predict estimated AD risk for other genotypes. A-F) miR-181d and miR-181c effects on probability of NCI, MCI, and AD. Primary effect is by miR-181d, expressed as modified by sex, brain region, and miR-181c levels. A) Temporal lobe (TL) Female. B) TL Male, C) Cerebellum (CB) Female, D) CB Male. E) PC Female, F) PC Male. G-I) Effects of APP and neprilysin mRNA levels on AD frequency. G) Effects on TL. H) Effects in PC, ages 76-85. I) Effects on PC ages 88-96.

### APP and neprilysin brain mRNA levels were associated with diagnosis probabilities of NCI, MCI and AD

When we modeled levels of APP and neprilysin mRNAs against diagnosis status of NCI, MCI, and AD in brain samples we found distinct effects depending on brain region. In temporal lobe we found that, unexpectedly, elevated APP mRNA tended to be associated with reduced probability of AD (Fig. 1G). However, this tendency reversed at lower levels of neprilysin mRNA. This may reflect deeper an interaction between APP and neprilysin mRNA levels, since it appeared that levels of APP and neprilysin mRNAs appeared negatively correlated in exploratory analysis (Supplemental Fig 2), and higher levels of APP would tend to be more common alongside lower levels of neprilysin. When the PCC was examined (Fig. 1H-I), increased APP mRNA levels associated with a greater probability of AD. This trend grew stronger with age. However, in PCC, levels of APP and neprilysin mRNA may be directly related, according to exploratory analysis (Supplemental Fig 2)

### SNPs near the *MIR181A1* and *MIR181B1* and neprilysin genes were associated with entorhinal cortex thickness

We modeled genotypes of rs111787496 (miR-181a and miR-181b associated) and rs111276850 (neprilysin associated) SNPs (Fig. 2A-B) against entorhinal cortical (EC) thickness. In our sample, rs111787496 only had AG and GG genotypes, while the neprilysin-associated SNP had AA, AG, and GG genotypes. We found that the rs111787496 GG genotype was associated with lower EC thickness (Fig. 2C-F). The neprilysin genotype interacted with sex. In female subjects, EC thickness increased as the number of minor “A” alleles decreases (AA→AG→GG) (Fig. 2C-D). In male subjects, EC thickness decreased as the number of minor alleles decreases (AA→AG→GG) (Fig. 2E-F).Genotype-associated changes were modest. Some of the data used in the preparation of figure 2 were obtained from the NIH-funded Alzheimer’s Disease Neuroimaging Initiative (ADNI) database. For up-to-date information, see adni.loni.usc.edu.

**Fig. 2.**
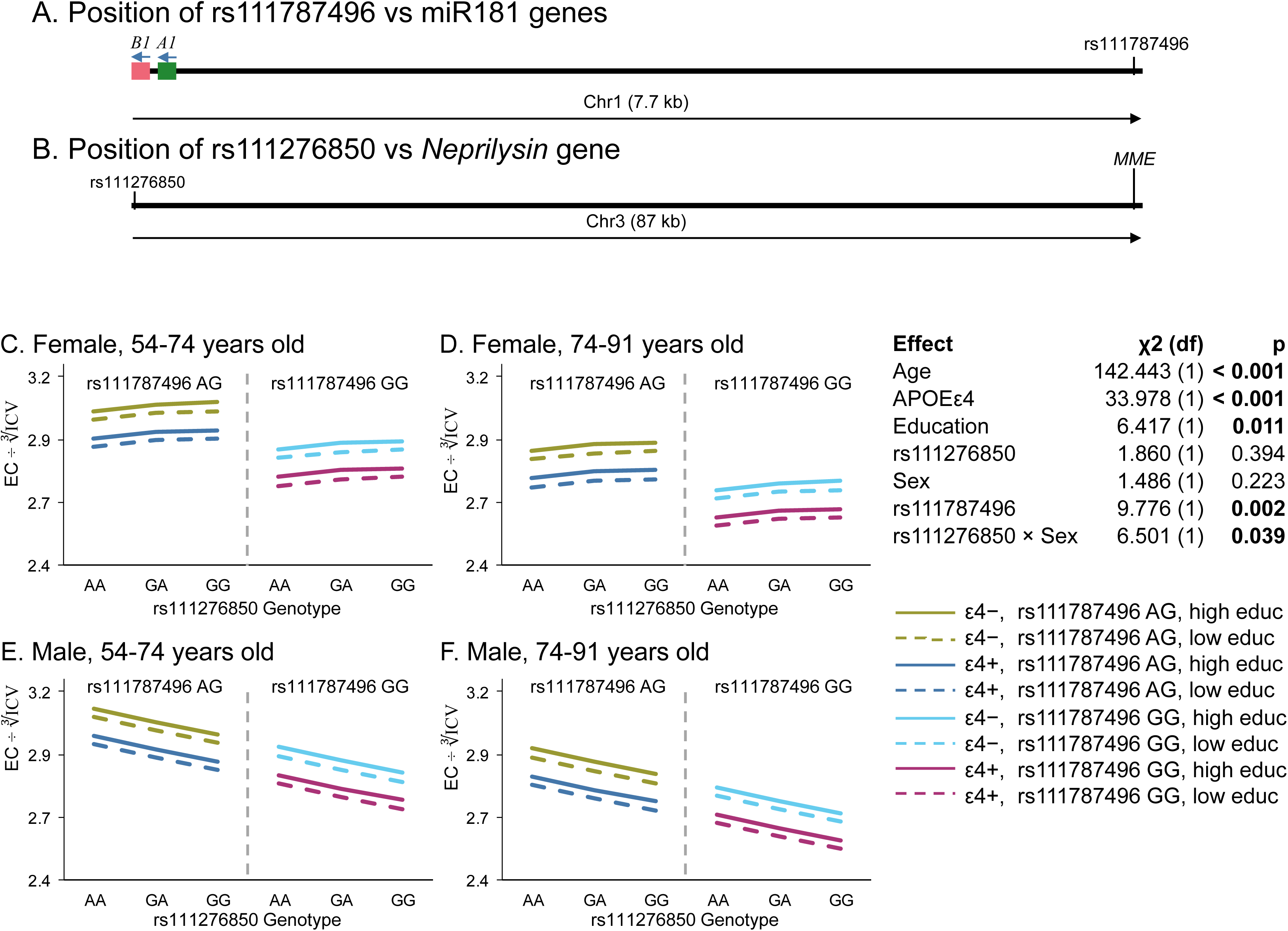
Genomic SNP effects on entorhinal cortex thickness. A) rs111787496 MIR181B1 and MIR181A1 genes on human chromosome 1. B) rs111276850 and the MME gene on chromosome 3. C-F) Effects of SNPs by sex and age. C) Female, 54-74. D) Female, 74-91. E) Male, 54-74, F) Male 74-91. We modeled genotypes of rs111787496 (miR-181a and miR-181b associated) and rs111276850 (neprilysin associated) SNPs against the entorhinal cortex (EC). In our samples, rs111787496 had only AG and GG genotypes, whereas the neprilysin-associated SNP had AA, GA, and GG genotypes. Each figure contains two halves, representing rs111787496 AG (left) and rs111787496 GG (right) as labeled in the top. The rs111276850 genotypes (AA, GA, and GA; x-axis) and EC thickness (y-axis) are shown. Our results suggest that the 111787496 GG genotype (right half) was associated with lower EC thickness than 111787496 AG genotype (left half) (Fig. 2C-F). The neprilysin genotype (y-axis) interacted with sex; specifically, in female subjects, EC thickness increased as the genotype progressed from AA to GA to GG (Fig. 2C-D). In male subjects, EC thickness decreased as genotype progressed from AA to GA to GG (Fig. 2E-F). For rs111787496 AG, the red straight line denotes e4 negative/ high education, and the broken lines e4 negative/low education. Deep blue denotes e4 positive (straight line: higher education; broken line: low). For rs111787496 GG, deep red and deep blue lines were used. It should be noted that any associated changes due to genotypes in either SNP were not large.

### Bioinformatics predict miR-181 sites in neprilysin mRNA 3’-UTR

Multiple online bioinformatics tools were consulted, including TargetScan, DIANA-microT, miRDB, RNA22, StarMir, and miRmap using different algorithms. By comparing and compiling data from these bioinformatics tools, we found two potential miR-181-5p binding sites on neprilysin 3’-UTR (Fig. 3A-C). The upstream seed sequence binding site is located at nucleotides 3011-3017 relative to the +1 TSS, while the downstream one at 4320 to 4326 nt relative to +1 TSS. The miR-181-5p seed sequence binding regions on neprilysin mRNA 3’-UTR are conserved in multiple mammal species, but not in mouse (Fig. 3B-C).

**Fig. 3.**
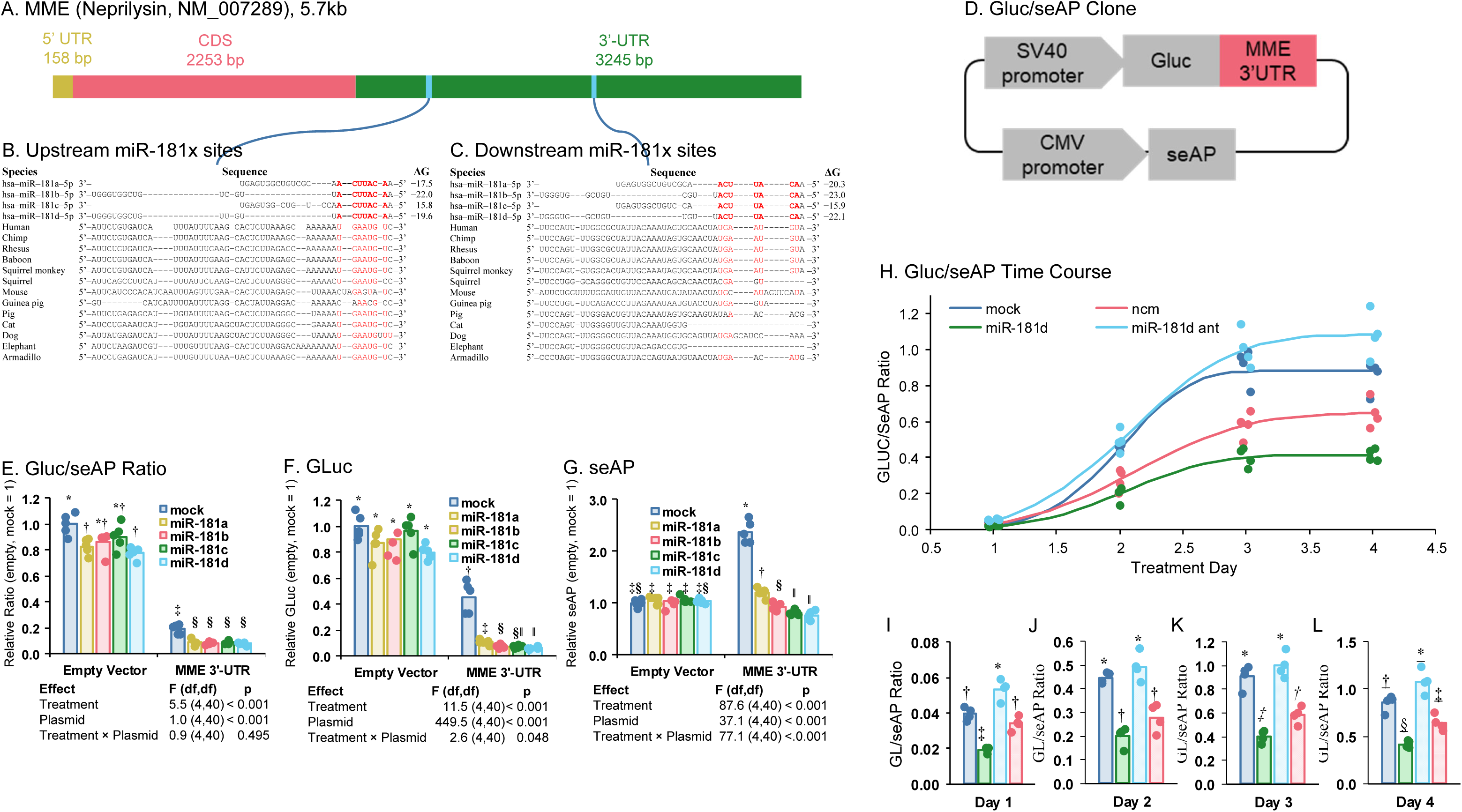
Presence of miR-181 target sites on the neprilysin mRNA, and the activity of neprilysin 3’-UTR altered by miR-181d. A) The neprilysin gene’s 3’-UTR is predicted to have B) an upstream and C) a downstream miR-181 binding site, both of which conserve the seed sequence strongly. D) Gluc/seAP (Secreted Alkaline Phosphatase and Gaussia Luciferase, two secreted luciferase) secreting reporter clone schematic with neprilysin 3’-UTR insert downstream of Gluc. The Gluc reading reflects neprilysin 3’-UTR activities, while seAP serving as an internal control. E) Ratio of Gluc/seAP levels representing neprilysin 3’-UTR activities in treated cells. F) Levels of corresponding Gluc in E. G) Levels of corresponding seAP in AE. H) Time course of GLUC/seAP secretion in cells under multiple treatments. I-L) GLUC/seAP secretion in each day separately.

### Four miR-181 forms targeted the neprilysin 3’-UTR sequence

We constructed a secreting Gluc/seAP reporter clone with the neprilysin 3’-UTR sequence inserted (Fig. 3D), as described herein. One set of cell cultures was co-transfected with mimics for miR-181a, b, c, and d. Then, media were sampled for Gluc and SeAP assays, the ratio of which was used as neprilysin 3’-UTR reporter readout. Each of the four miR-181 forms reduced reporter output (Fig. 3E-G). However, there was also some interaction of the miRNA mimics and the empty vector, and the neprilysin 3’-UTR on its own significantly altered reporter expression. We also treated this clone with miR-181d mimic, an antagomir to miR-181d, and a noncomplementary oligo and collected media over 4 days. Gluc/seAP levels were measured and analyzed with nonlinear regression, using the Weibull 2.3 model. This model has three coefficients, specifically asymptote (maximum signal), steepness (slope of ascending portion) and inflection (in days). We found that treatment with miR-181d significantly reduced asymptotic maximum and that treatment with antagomir significantly increased maximum signal vs. mock (Fig. 3H-L). In addition, miR-181d treatment significantly decreased the curve inflection point and steepness. Overall, compared to the mock curve, Gluc/seAP ratios for miR-181d treated cells followed a shallower curve that rose more slowly and never reached as a high level. On the other hand, blockade of endogenous miR-181d by antagomir produced a response that initially rose more slowly than untreated but eventually resulted in the highest levels of Gluc/seAP ratio, suggesting detectable background activity of miR-181d levels in the cells.

### miR-181 reduced neprilysin mRNA and protein

All four members of the miR-181-5p family reduced neprilysin protein, but not Aβ related proteins APP or ECE1 (Fig. 4A-B). BACE1 protein was significantly up-regulated by miR-181d. The regulation could be through direct miRNA binding on BACE1 mRNA or indirect effects potentially through Aβ as a transcription factor (*15, 16*). To further confirm the role of miR-181d-5p on neprilysin regulation, neprilysin mRNA and protein levels were determined by real-time qPCR and western blotting (Fig. 5A-B). miR-181d-5p significantly reduced neprilysin mRNA and protein levels as compared to mock transfection (Fig. 5B-C), consistent with miR-181d reducing neprilysin expression at least partly through reduced neprilysin mRNA abundance. miR-216, another predicted neprilysin regulator, reduced neprilysin mRNA but did not reproduce the full miR-181d protein-level pattern..

**Fig. 4.**
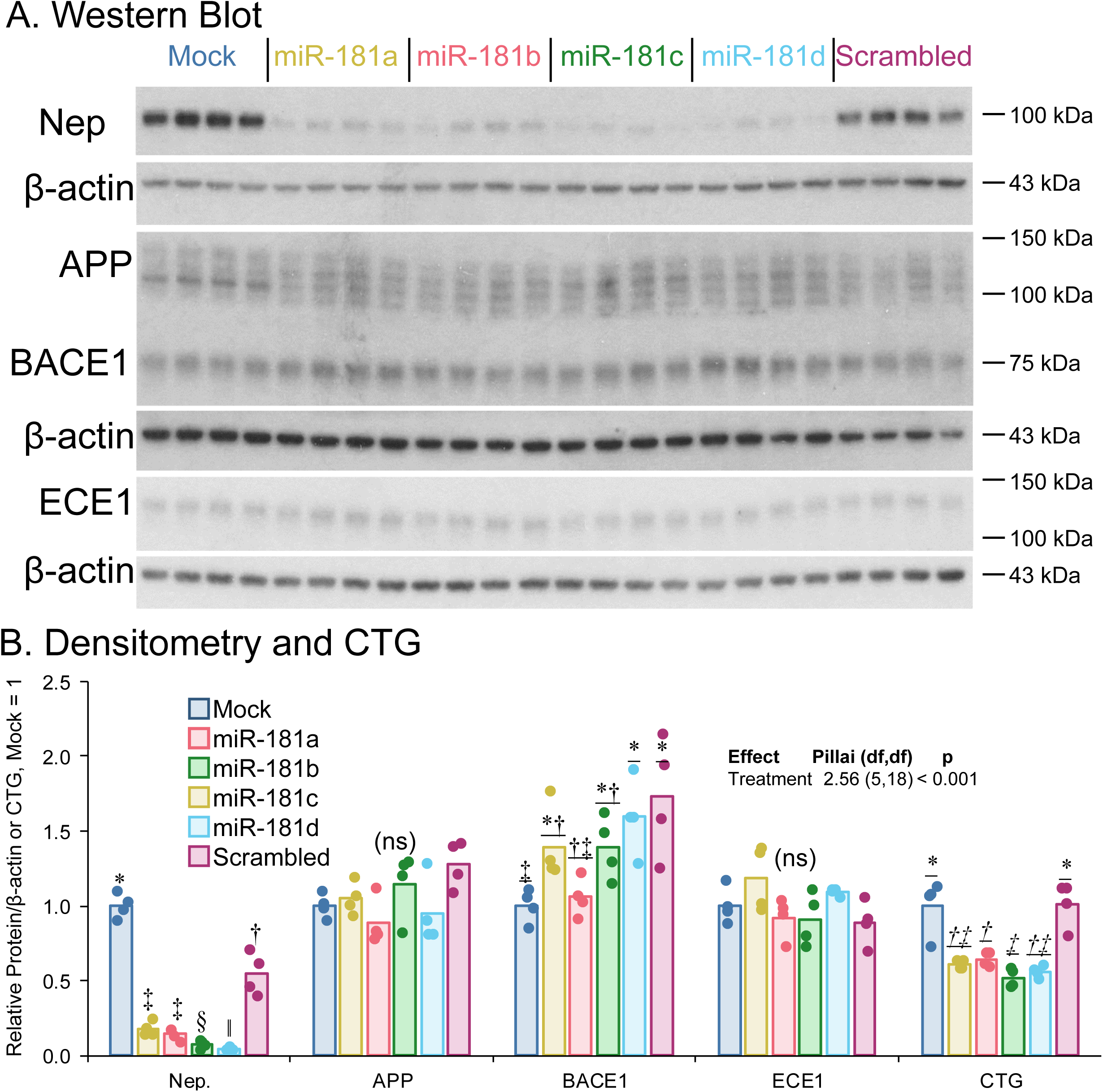
MiR-181a, b, c, d reduce endogenous neprilysin protein in differentiated SK-N-SH cells. western blotting of miR-181d effects on MME level and activity and on Aβ. SK-N-SH cells were either mock-transfected or transfected with 100nM miRNA mimics or scrambled miRNA mimics. The scrambled miRNA mimic is a non-targeting control miRNA mimic, with no known impact on miRNA functions. A) Western blotting of Neprilysin, APP, BACE1 and ECE1 proteins. There were three gels separately probing neprilysin, APP together with BACE1 and ECE1. Each of the three gels have their corresponding β-actin as loading controls. B) Densitometric quantification of the target proteins and corresponding β-actin on the same blot as well as CellTiter-Glo (CTG) cell viability assay.

**Fig. 5.**
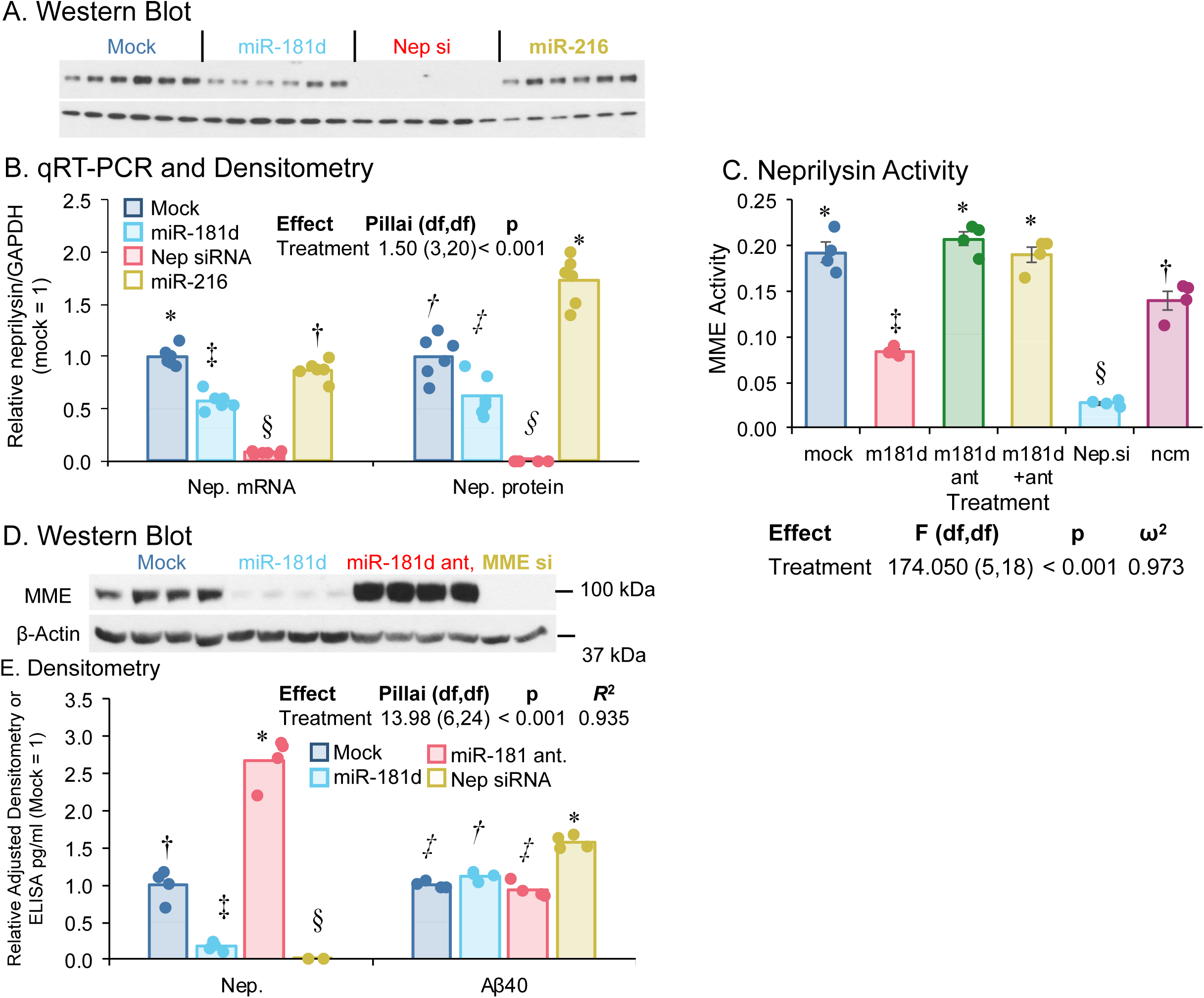
MiR-181d reduces endogenous neprilysin protein in differentiated SK-N-SH cells. MiR-181d mimics, neprilysin siRNA and miR-216 mimics, another miRNA predicted to target neprilysin were transfected into differentiated SK-N-SH cells, from which, both protein and mRNA were extracted. A) Western blot of cell extracts. B) qRT-PCR of MME mRNA and Densitometric quantification of protein levels. C) Neprilysin Activity. D) Western blot of neprilysin in transfected SK-N-SH cells. E) Densitometry of neprilysin blot and levels of Aβ 1-40 ELISA levels assayed from the conditioned media of transfected cells.

### miR-181d reduced neprilysin enzyme activity and increased Aβ levels

miR-181d transfection significantly down-regulated neprilysin protein and enzyme activity (Fig. 5B-C). Treatment with its specific antisense inhibitor, antagomir alone or in combination reversed its effects. Since neprilysin is a well-established Aβ degrading enzyme, we determined the levels of Aβ 1-40 in the conditioned media (Fig. 5E). Extracellular Aβ1-40 levels showed a modest but statistically significant increase after miR-181d treatment.. Treatment with neprilysin siRNA increased Aβ levels as a positive control. However, while treatment with miR-181d reduced neprilysin protein levels in the same cells, miR-216 treatment did not alter neprilysin protein levels. Notably, miR-181d-5p antagomir transfection alone increased neprilysin protein levels (Fig. 5D-E), suggesting the active and potent regulation of neprilysin by endogenous miR-181d-5p in cells.

### Bioinformatics prediction revealed that miR-181 sites in MAPT mRNA CDS and 3’-UTR

Bioinformatics prediction (Fig. 6A-D) suggested that miR-181-5p binds to MAPT coding sequence (CDS) and 3’-UTR (NM_005910). Two binding sites were located at MAPT 3’-UTR 4933-4957 and 5622-5640 nt relative the +1 transcription start site (TSS), while another binding site located on MAPT CDS 984-1007 nt relative the +1 TSS.

**Fig. 6.**
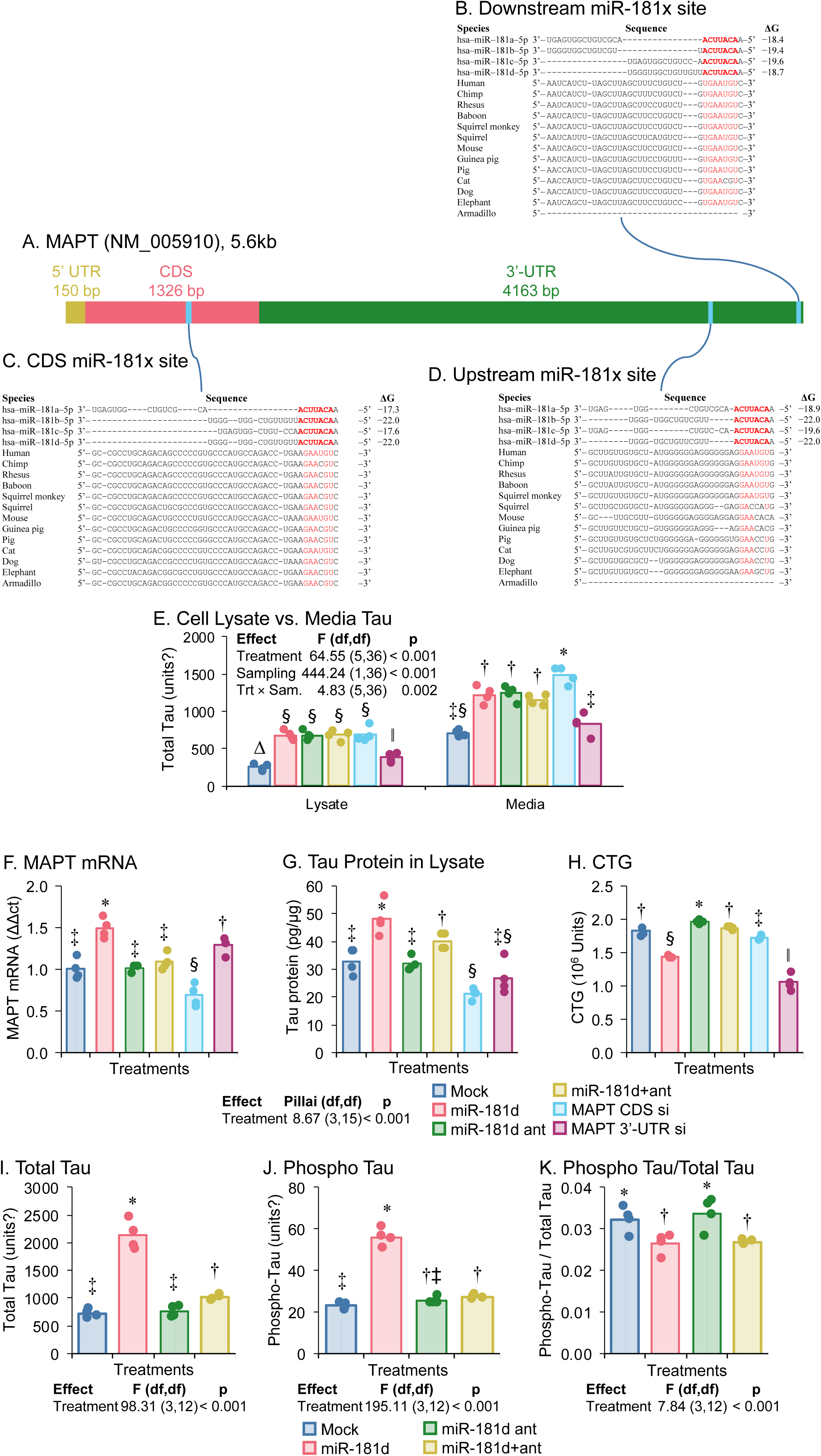
MiR-181d increased tau mRNA and protein. A) Sequence of MAPT mRNA. B-D) Bioinformatic prediction of miR-181d binding sites on MAPT mRNA CDS and 3’-UTR. SK-N-SH cells were transfected with miR-181 family or miR-181d along with its antagomir. The cell lysate or conditioned media were subjected to tau ELISA assay. E) Total tau protein levels by ELISA of cell lysate and conditioned media. F) MAPT mRNA real-time qPCR quantification. G) Tau protein ELISA of cell lysate. H) CTG. I-K) Total tau and phosphor-tau (pT231) ELISA of cell lysate.

### miR-181 transfection increased tau protein and mRNA levels, without altering UTR reporter activity

Four miR-181 family members miR-181a, b, c and d all significantly increased tau protein levels, both total tau protein in the cell lysate and secreted tau protein in conditioned media (Fig. 6E). As miRNAs typically bind to 3’-UTR, the RISC complex represses target mRNA expression. To further confirm this result, tau mRNA levels were determined in cells transfected with miR-181d, its antisense specific antagomirs alone or in combination. miR-181d transfection significantly increased both tau mRNA (Fig. 6F) and protein (Fig. 6G) levels. miR-181d antagomirs reversed miR-181d effects on tau, confirming target binding specificity. Two MAPT siRNAs were also applied here, one targeting CDS while another targeting 3’-UTR. The CDS-targeting siRNA reduced tau protein and mRNA validating assays specificity. miR-181d application further associated with reduced cell viability as measured by CTG (Fig. 6H). To pinpoint the binding sites of miR-181d on MAPT mRNA, the MAPT 3’ and 5’-UTR reporter clones were generated and assayed. However, miR-181d transfection did not alter either reporter, indicating and untested site or potential indirect effects on MAPT mRNA by miR-181d (Supplemental Fig. 3).

### miR-181d increased phosphorylated tau protein levels

Since total tau protein was up-regulated by miR-181d transfection, we tested whether the phosphorylated tau levels were altered. The phosphorylated tau at site Thr231, which was found to be an early phosphorylation site, was accordingly increased as well. The ratio of phospho-tau vs total tau did was significantly increased, suggesting that tau related kinases and phosphatases were altered by miR-181d transfection (Fig. 6I-K).

### RNA-seq identified miR-181d-regulated mRNAs enriched for AD- and neurodegeneration-associated pathway annotations

miR-181d, antagomir, and mock transfections in differentiated neuroblastoma cells were subjected to RNA sequencing. The results were filtered and used for network analysis as described herein. Expression of 3,120 mRNAs was altered by miR-181d vs. mock transfection comparison, 3,736 were altered in the antagomir versus mock comparison, and 304 were altered in the miR-181d versus antagomir comparison. (Supplemental Data). It should be noted that a significant contrast of miR-181d vs antagomir would mean that the interaction was strongly asymmetrical, in that the net outcome was not “reset to untreated” and revealed a failure of antagomir/miR-181d interaction. Using pathway-guided filtering for exploratory network analysis, 414 genes had significant contrasts of miR-181d vs mock (Fig. 7A), 51 had significant contrasts of miR-181d vs. antagomir (Fig. 7B), and 509 had significant contrasts of antagomir vs. mock (Fig. 7C). These analyses were descriptive and hypothesis-generating. Overall net effects, summarized by examining net treatment effect vs. total effect magnitude and filtered by opposition index ≥7 (which corresponded to ≥0.49 on a 0-1 scale), resulted in 57 “seed” genes, to which MME and MAPT were added for network analysis. Hippocampus network (Fig. 7E) was very tightly interlocked and densely interconnected. The network included several genes annotated as directly involved in Aβ, p-tau, or α-synuclein aggregation or clearance (Supplemental Table 1), revealed several among the seeds and those pulled out by network analysis, highly interconnected. Clustered KEGG analysis of this network indicated a tightly overlapping set of pathways (Fig. 7F) that are also functionally related, all represented among miR-181d-responsive genes in this exploratory analysis. When network analysis was performed on the frontal cortex PPI database (Fig. 7G), a similar tightly interlinked network appeared. The clustered KEGG analysis (Fig. 7H) produced a larger clusters. For both hippocampus and frontal cortex networks, the KEGG cluster containing the Alzheimer’s disease annotation separated from other pathway clusters in this exploratory analysis (Supplemental Table 2, Supplemental Figs 4-5)

**Fig. 7.**
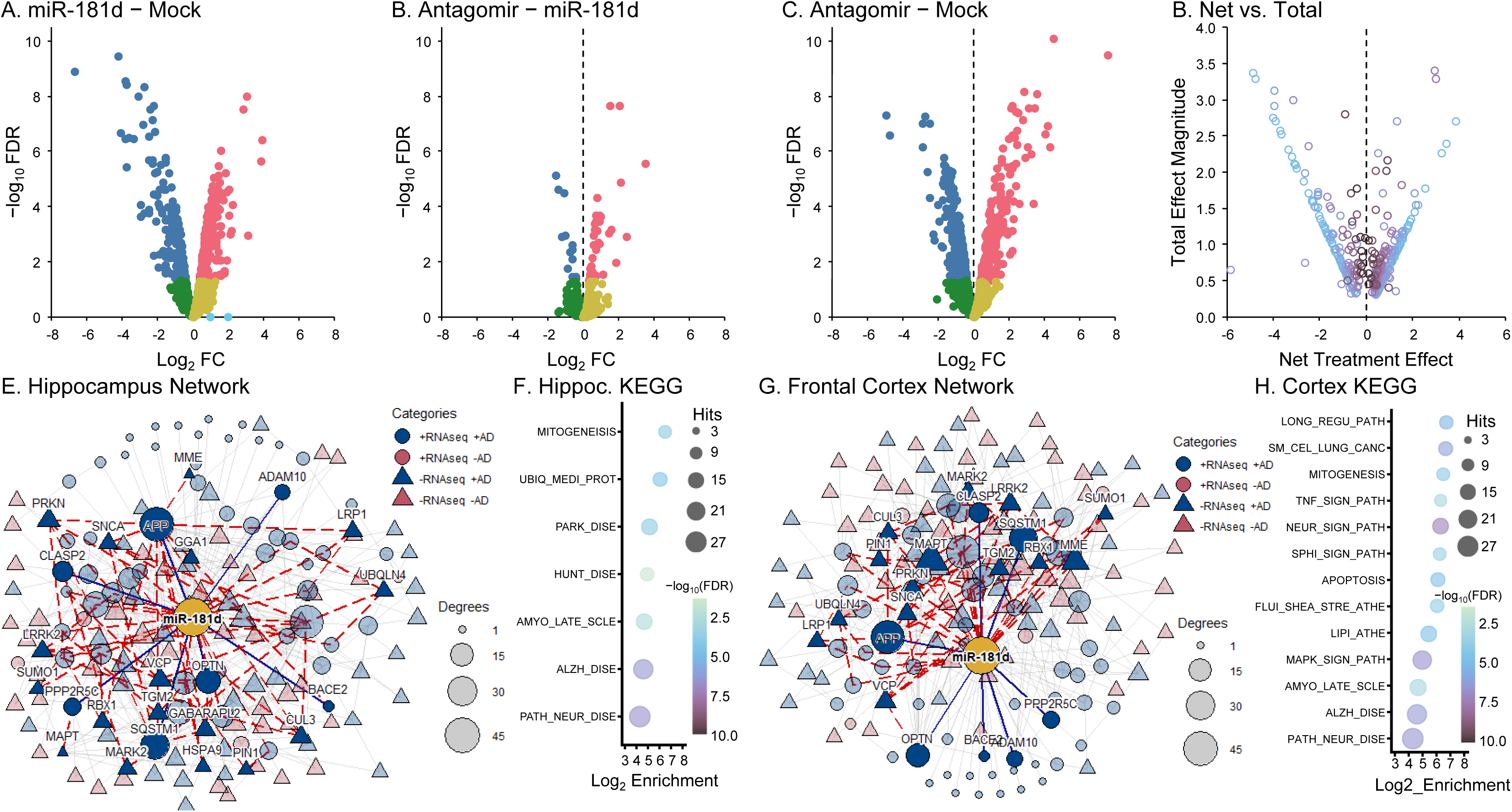
Targeted RNA-seq to network pathway analysis based on miR-181d activity. RNA-seq was carried out and analyzed as described herein. RNA-seq outcomes are A) Volcano plot of miR-181d vs. mock contrast (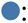 decreased, FDR ≤ 0.05; 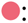 increased, FDR ≤ 0.05; 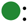 decreased, FDR > 0.05; 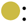 increased, FDR > 0.05). B) Volcano plot of antagomir vs. miR-181d contrast. C) Volcano plot if antagomir vs. mock contrast. D) “pseudo-volcano” plot of composite indices (total effect magnitude vs net treatment effect), color coded by opposition index strength clusters (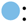 0; 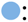 >0 to <0.10; 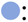 0.10 to <0.19; 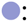 0.19 to <0.27; 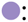 0.27 to <0.34; 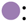 0.34 to <0.41; 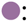 0.41 to <0.49; 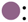 0.49 to <0.58; 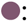 0.58 to <0.68; 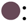 0.68 to <0.81; 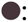 0.81+). E) Hippocampus network from opposition index screened genes. “Degrees” refer to the number of connections a node has to other nodes. F) AD-containing KEGG pathway cluster associated with hippocampus network. G) Frontal cortex network from screened genes. H) Frontal Cortex KEGG AD-associated cluster.

### Levels of miR-181d, MAPT mRNA, and neprilysin mRNA differed by cell differentiation

When we differentiated iPSC cells into neuropgenator cells, neurons, and astrocytes (Fig. 8 B-D), we found that levels of each differed by cell type and were dissimilar.

**Fig. 8.**
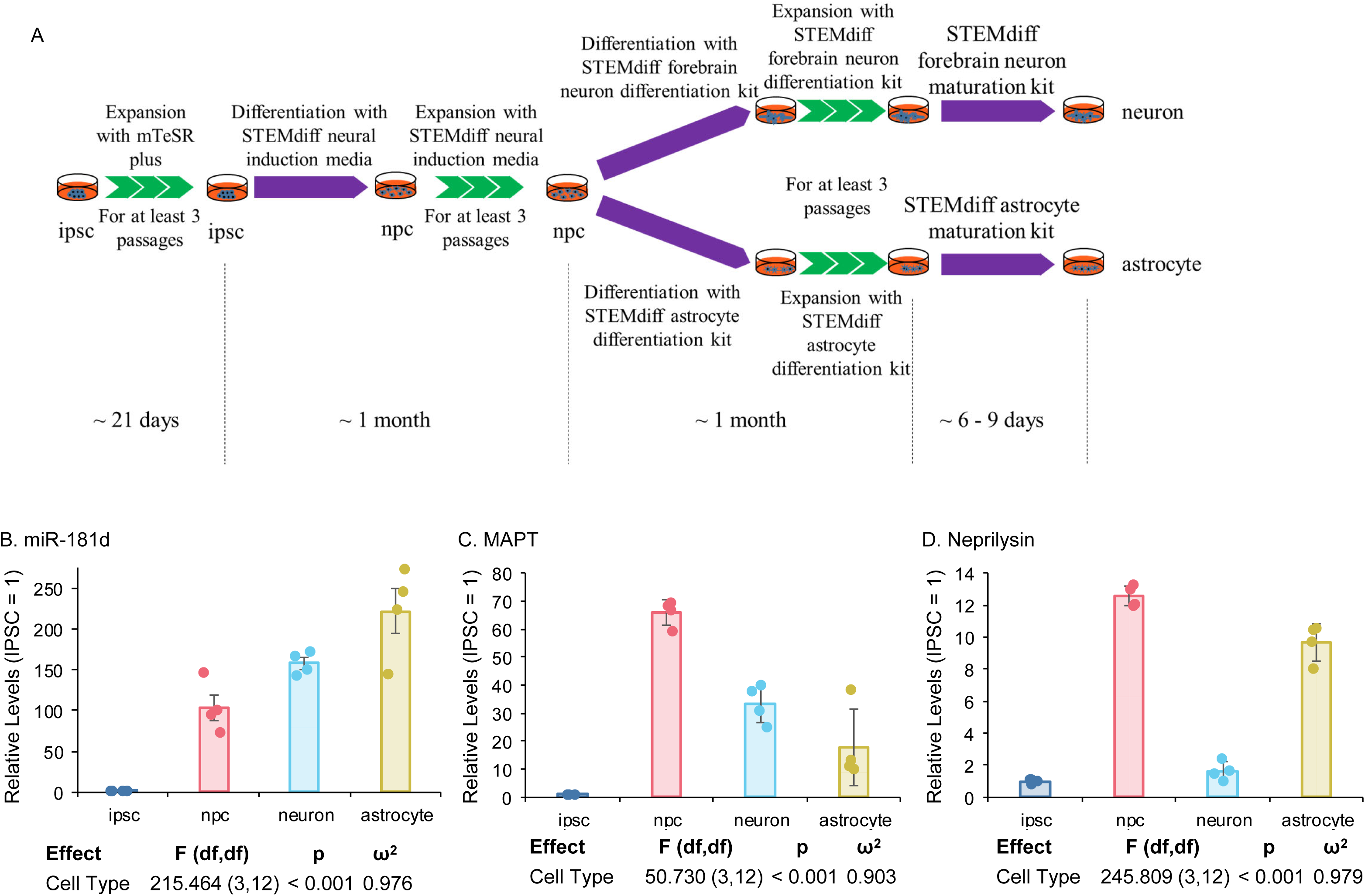
IPSC cell neuronal differentiation and miR-181d, MAPT, neprilysin levels. A) ipsc cells were differentiated as illustrated. B) miR-181d levels in different stages of ipsc differentiation. C) MAPT mRNA levels. D) Neprilysin mRNA levels.

## Discussion

In this study, we found that miR-181d-5p levels in human brain samples from temporal lobe, cerebellum and posterior cingulate cortex were significantly associated with AD diagnosis probability. Overall, as miR-181d levels increased, probability of AD diagnosis increased. This trend was particularly strong in female subjects than in male subjects. However in the PCC region of male patients, miR-181d is negatively associated with AD probability. Presence of the APOEε4 allele also increased likelihood of AD diagnosis, independent of miR-181d levels.

The miR-181 family contains miR-181a-5p, miR-181b-5p, miR-181c-5p and miR-181d-5p, which are derived from six different sequences of three chromosomes (chromosomes 1, 9 and 19). The four miR-181-5p species share the same seed sequence and hence share a great similarity in target binding. It is worth noting that besides these four miR-181-5p, there are six other miR-181-3p generated together with their corresponding miR-181-5p segments. miR-181, as a big miRNA family, is involved in multiple biological and pathological processes (*17*).

Many therapeutic strategies have focused on removing Aβ, reducing Aβ generation, or preventing Aβ aggregation on removing Aβ by active or passive immunotherapies, reducing Aβ generation by β-secretase or γ-secretase inhibitors, or preventing Aβ aggregation. However, Aβ proteolytic clearance pathways have been less extensively explored. Since neprilysin is a key Aβ-degrading enzyme, increased miR-181d could reduce neprilysin-dependent Aβ clearance in AD-relevant cellular contexts (*4, 18*). The magnitude of Aβ1-40 increase in miR-181d-transfected neuronal cell cultures was limited, possibly because of the short incubation time and low endogenous Aβ levels.

Neprilysin expression levels are highest in kidney. Thus, there is a hypothesis that Aβ clearance is accomplished primarily in kidney and other peripheral organs expressing neprilysin but not brain. Neprilysin gene transfer in skeletal muscle of AD mice significantly reduces Aβ burden in the brains (*19*).

miR-181 treatment increased tau mRNA and protein. The exact mechanism remains unresolved. We cloned both MAPT 5’ and 3’-UTR luciferase clones and co-transfected with miR-181 in differentiated neuroblastoma cells. However, miR-181 did not alter either 5′- or 3′-UTR reporter activity, indicating that direct MAPT targeting through the tested UTRs was not demonstrated. One predicted binding site lies in the MAPT CDS, but this site was not directly tested here. Alternatively, miR-181 may indirectly alter MAPT expression through upstream transcriptional regulators. Although hyperphosphorylated tau protein is the major component of neurofibrillary tangles, tau protein under physiological circumstances is a microtubule binding protein, promoting microtubule stability and influencing axonal transport (*20*). The knockdown of tau protein inhibits neurite outgrowth in primary neuronal culture (*21*).

In addition to neurodegenerative diseases, miR-181 also plays a critical role in autism spectrum disorder (ASD). miR-181 levels are differentially expressed in the brain and serum of ASD patients (*22, 23*). miR-181-regulated genes are enriched in neurite and synapse developmental processes, leading to increased synaptogenesis and altered neurite outgrowth (*24*). miR-181 plays a critical role in synaptogenesis early in development and a compensatory role in neurodegenerative conditions of the brain later in life; consequently, excessive neuronal projection and synaptic maintenance under a metabolic-stress state in pre-AD ultimately lead to neurodegeneration. In that way, miR181d could play a positive role.

miR-181 is also involved in various neurological and neurodegenerative diseases, such as stroke (*25*), Parkinson’s disease (*26*), multiple sclerosis (*27*), cerebral ischemia (*28*), and amyotrophic lateral sclerosis (*29*). Circulating miR-181 could serve as a prognostic biomarker for amyotrophic lateral sclerosis (*29, 30*). Conversely, miR-181 promotes progression of or resistance to multiple cancers including bleomycin-induced skin fibrosis, hepatocellular carcinoma and glioblastoma (*31–34*). miR-181 also regulates expression of genes involved in apoptosis and autophagy, including Bcl-2, ATG5, and ATG7 (*35–37*). miR-181 overexpression in astrocytes leads to increased secretion of the anti-inflammatory cytokine IL-10. By contrast, miR-181 knockout is pro-inflammatory, increasing the production of Il-1β, IL-6, IL-8 and TNF-α (*38*). Likewise, specific miRNAs can regulate pathways that participate in both early-life conditions, such as ASD (*22*), and late-life disorders, such as ADRDs.

At cellular level, human neuroblastoma cells are also used as proxies for human neurons. These cells are “differentiated” into neuron-like cells to test neuroprotective strategies for AD like condition and cell survival (*39*). Recent research has identified specific transcription factors, such as DMRT2, that play roles in both the development of the cingulate cortex and the proliferation of neuroblastoma cells (*40*). Using differentiated neuronal cells, we showed that miR-181 treatment increased levels of synaptic protein markers. miR-181d increased doublecortin and SNAP-25 mRNA, by RNA-seq, in addition to Western blot. PSD-95 (post-synaptic density protein) showed a trend towards increased expression. (Wang et al-data not shown). Therefore, the present work unveiling the role of miR-181d in neprilysin, tau, and synaptic proteins assumes a great significance in both ASD and ADRDs.

In summary, our data shows the miR-181d dysregulation is associated with alterations in multiple AD-relevant molecular pathways that play important roles in AD etiology. miR-181d is associated with altered probability of an AD diagnosis. A SNP in proximity to *MIR181A1*/*MIR181B1* is associated with altered entorhinal cortex thickness. miR-181d targets neprilysin mRNA 3’-UTR, reducing neprilysin mRNA, protein and enzymatic activities. Consistent with reduced neprilysin activity, miR-181d modestly increased extracellular Aβ1-40 in conditioned media. Moreover, miR-181d increased tau protein and phospho-tau protein levels, with a significant, if small, reduction in phospho-tau to tau protein ratio. RNA sequencing followed by exploratory enrichment analysis identified miR-181d-regulated mRNAs enriched for AD- and neurodegeneration-associated pathway annotations.

We propose a partial model relating miR-181 family members to AD (Fig. 9). Specifically, miR-181-5p members, which could include miR-181a, miR-181b (SNP data) or miR-181d (protein and transcriptomics) would influence translation (if not necessarily levels) of neprilysin and tau. Specifically, diminution of neprilysin, which impairs Aβ clearance and elevation of tau, would contribute to plaque and tangle formation, respectively. Other AD-related proteins may also be perturbed by changes in miR-181. The collective effect of these changes leads to greater downstream risk of AD. Altogether, our results suggest that neprilysin diminution impairs Aβ clearance and elevates tau, contributing to AD pathology. RNA sequencing identified miR-181d-responsive neurodegenerative pathways associated with neurodegeneration. Diminution of neprilysin, which impairs Aβ clearance and elevation of tau, would contribute to plaque and tangle formation, respectively. The collective effect of these changes leads to greater downstream risk of AD. These findings identify miR-181 as a regulator of AD-relevant amyloid and tau pathways, and potentially provide novel targets for AD.

**Fig. 9.**
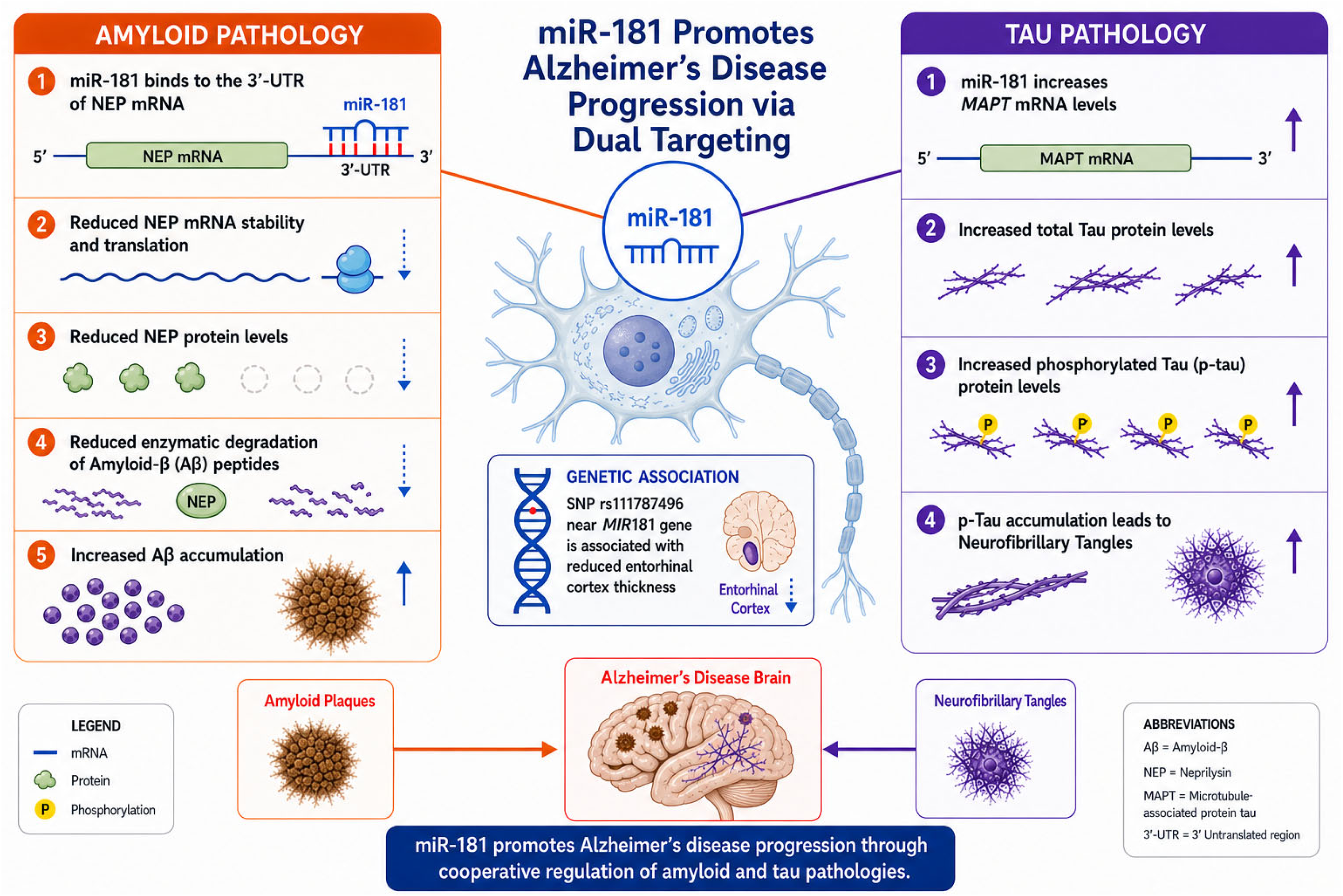
Directional model summary of implications of miR-181d in Alzheimer’s disease. MiR-181 reduced neprilysin mRNA and protein levels, thus increasing Aβ accumulation. Furthermore, miR-181 increased tau mRNA and protein as well as phosphor-tau protein levels, contributing to AD progression. miR-181d levels are associated with altered AD probability. A SNP near miR-181 gene is associated with reduced entorhinal cortex thickness.

## Materials and Methods

### Identification of putative miR-181-5p binding sites on neprilysin mRNA 3’–UTR

Multiple bioinformatic prediction tools with different algorithms were applied to find the binding site of miR-181-5p in neprilysin mRNA 3’–UTR, including TargetScan, DIANA-microT, miRDB, RNA22, StarMir and miRmap (*41–48*).

### Cell culture

Human glioblastoma cells (U373), HeLa cells, neuroblastoma cells (SK-N-SH), and microglia cells (HMC3) were obtained from the American Type Culture Collection (ATCC). Cells were grown in Eagle’s modified minimum essential media (EMEM) (Corning) containing 10% fetal bovine serum (FBS) (Atlanta Biologicals) and penicillin/streptomycin solution (Corning) at 37 °C in 5% CO_2_ humid incubators. Human neuronal cultures were prepared by differentiating human neuroblastoma (SK-N-SH) cell line at 50% confluence in the presence of all-trans retinoic acid (ATRA, Sigma) for 7 days in low serum medium (1% FBS).

### Transfection of nucleic acids

Transfections were performed when cells reached around 80% confluence. Culture media were replaced with Opti-MEM media (Gibco) with 1% FBS and antibiotics were omitted from transfection media. For miRNAs and siRNAs transfection, Lipofectamine RNAiMax (Invitrogen) was applied 2μ l per well in a 24 well plate format. 75nM miRNA (Ambion) or 50nM siRNA (Santa Cruz Biotechnology) were premixed with RNAiMax according to its protocol. Mixes were added into each well and kept for 72 hours or otherwise indicated in figure legends. For co-transfection of plasmid DNA with miRNA, plasmids (Genecopia) were added at 100ng/well in a 96-well plate or 400ng/well in 24 plates. RNAiMax and miRNA were scaled down or up accordingly.

### Gluc/seAP reporter clone assay

The pEZX-MT05 vector (GeneCopeia) has both a *Gaussia* luciferase (GLuc) and secreted alkaline phosphatase (seAP) fused in-line with each other, with a multiple cloning region immediately downstream of the GLuc coding sequence (Fig. 2D). This permits seAP to act as a strong control for transfection efficiency while GLuc levels can be modified via a 3’-UTR insert. Neprilysin 3’-UTR was cloned into this vector and co-transfected along with microRNA mimics into ATRA-differentiated SK-N-SH (NBRA). Conditioned media were harvested and subject to luminometer measurements of GLuc and seAP activity (Fig. 2E-H). We have succeeded miRNA transfection in poorly transfectable neuronal cultures by optimizing several parameters (data not shown). Transfection in neuronal cultures was performed without the presence of ATRA.

### Cell lysing

After washing with PBS, cells were lyses on-plate with vigorous shaking using 100 RIPA buffer (Thermo Scientific) containing 1X Halt Protease Inhibitor Cocktail (Thermo Scientific). Protein concentration was determined by BCA assay (Thermo Scientific) according to the manufacturer’s protocol and then Laemmli sample buffer (LSB) (Bio-Rad) was added to each tube of lysate. Lysate and LSB mixes were boiled for 10 minutes and cooled down on ice or kept in freezers for further studies.

### Human induced pluripotent stem cells (iPSCs) culture

A human iPSC cell line was obtained from Coriell Institute for Medical Research (#AG25370, derived from the skin fibroblast of a white 81-year-old German female familial AD patient harboring PSEN2 mutation). IPSC cells were cultured as previously described(*49*). Briefly, cells were maintained and expanded with mTeSR plus media (STEMCELL Technologies) and then differentiated into neural progenitor cells (NPCs) with STEMdiff™ SMADi neural induction kit using a monolayer culture protocol. NPC cells were further differentiated into forebrain neurons and astrocytes with STEMdiff™ neuron differentiation kit and maturation kit, STEMdiff™ astrocyte differentiation kit and maturation kit respectively. Protein and RNA were harvested at different stages for further analysis.

### SDS–polyacrylamide gel electrophoresis (SDS–PAGE) and Western immunoblotting

An equal protein amount of lysate was loaded onto 26 lane BisTris XT denaturing 4-12% polyacrylamide gels and run with XT MOPS buffer (Bio-Rad). Proteins were separated with SDS-PAGE and then transferred overnight onto PVDF membranes (Bio-Rad). Membranes were stained with 0.1% Ponceau S solution to confirm transfer success. After three times of washing with TBST, membranes were incubated with 5% nonfat milk in TBST for 1 hour at room temperature. Primary antibodies were incubated at either room temperature for 3 hours or 4 °C overnight. Goat anti-rabbit or mouse secondary antibodies (Invitrogen) were applied for 1 hour at room temperature. Protein bands were visualized using auto radiographic films (Dot Scientific) and ECL buffer (Thermo Scientific). Films were scanned for densitometry analysis. Primary antibodies were used to probe neprilysin (Abcam ab208778), APP (Millipore MAB348), BACE1 (Cell Signaling D10E5), ECE1 (Abcam ab71829) and β-actin (Sigma A-5441)

### RNA isolation and real-time quantitative PCR

Total RNA was isolated with mirVana miRNA isolation kit (Invitrogen) following its protocol. RNA concentration and quality were assessed by Nanodrop instrument. Equal amount of RNA per sample was reverse transcribed with High-Capacity RNA-to-cDNA kit (Applied Biosystems) for mRNA quantification. cDNA was subjected to real-time qPCR analysis. Relative quantification was achieved by delta delta Ct normalization with the geometric means of housekeeping genes GAPDH and β-actin.

### Enzyme Labeled Immunosorbent Assay (ELISA)

Specific neprilysin ELISA (RnD) was performed following the kit protocol. Briefly, plates were coated with a capture antibody targeting neprilysin. After a blocking step, lysate, detection antibody, and HRP-conjugated to biotin-streptavidin steps for visualization were performed. To reduce non-specific results, we tested each sample both with and without the addition of the detection antibody. The ODs were then calculated as OD with detection antibody – OD without detection antibody. The resultant OD values were normalized to the mean of the control values and exhibited as shown. Likewise, levels of Aβ40 and Aβ (1-42) were measured independently by specific and sensitive ELISA as per the manufacturer’s protocol (IBL).

Neprilysin enzyme activity assay. Neprilysin activity was determined with a specific neprilysin activity assay kit (Biovision) following the manufacturer’s directions. Briefly, miRNA-or siRNA-transfected cells were lysed with assay buffer containing protease inhibitors. Three to five micrograms of each lysate was added to 96 well opaque plates. Neprilysin substrates were applied, and the solutions were mixed. The cleavage of neprilysin-specific synthetic substrates released free fluorophores, which are activated at 330 nm and emit at 430 nm. The plates were read using a fluorescence plate reader. Readings were taken every 5 minutes for 2 hours. The kinetics represented neprilysin enzyme activity.

### Human Brain Specimens and Processing

Postmortem brain tissue samples from three regions, the temporal lobe, cerebellum, and posterior cingulate cortex (PCC) used in this study has recently been described by us (*49*). Samples were procured by our collaborators Drs. Nelson and Counts. Samples were frozen in liquid nitrogen, powdered with a hammer, and subsequently lysed using M-Per buffer containing a protease inhibitor cocktail (Roche). After centrifugation, supernatants were boiled in Laemmli sample buffer and separated on SDS-polyacrylamide electrophoresis. A mix was prepared that combined small volumes from all samples to prepare a standard curve.

### SNPs in proximity to the *MIR181A1* and *MIR181B1* and neprilysin genes in the ADNI cohort

The ADNI initial phase (ADNI-1) was launched in 2003 to test whether serial magnetic resonance imaging (MRI), position emission tomography (PET), other biological markers, and clinical and neuropsychological assessment could be combined to measure the progression of mild cognitive impairment (MCI) and early AD 40, 41. ADNI-1 has been extended in subsequent phases (ADNI-GO, ADNI-2, ADNI-3, and ADNI-4) for follow-up of existing participants and additional new enrollments. More information about ADNI can be found at https://adni.loni.usc.edu. Demographic information, pre-processed neuroimaging scans, APOE and whole-genome genotyping data, neuropsychological test scores, and clinical information are publicly available from the ADNI LONI data repository.”

### RNA-seq Analysis

RNA-seq analysis began from a gene-by-sample matrix of gene identifiers, treatment identifiers, and gene-level read counts provided by the sequencing core facility; upstream processing performed by the IUSM genomics core. Samples were given one of three treatments: Mock, miR-181d, and antagomir, with four replicates per treatment. Gene-level counts were analyzed using an overdispersed count generalized linear model. Dispersion diagnostics indicated that the standard edgeR estimation a quadratic mean–variance component was excessive (i.e., the fitted mean–variance inflation appeared larger than warranted by the data), so we used a Gamma–Poisson GLM framework as implemented in glmGamPoi, with gene-wise overdispersion estimation and likelihood-based model fitting/testing appropriate for count data.

To characterize and prioritize miRNA-antagomir behavior for downstream network analysis, three empirical, descriptive composite indices were computed from two signed, model-estimated effects (b1: miR-181d−Mock and b2: Antagomir–miR-181d). These indices were used only for empirical ranking and pattern characterization, not as formal statistical contrasts derived by algebraic substitution: net treatment effect: b1 + b2; total effect magnitude: |b1| + |b2|; and opposition index 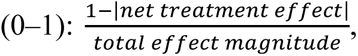 when total effect magnitude > 0; values near 1 indicate near-complete cancellation (opposing directions with similar magnitude), and values near 0 indicate reinforcement in the same direction.

Opposition index values of 0 were treated as a separate “zero-cancellation” category and excluded from further analysis, and opposition index > 0 values were clustered using pamk (partitioning around medoids with automatic k selection) into 10 clusters to create an ordinal binning of cancellation strength for ranking and selection (Supplemental Table 1). Genes were subsequently ranked according to this opposition index clustering structure; downstream selection used a truncation threshold of cluster 7, with MME and MAPT additionally included by design.

The Enrichr website was consulted using the full gene list to probe the Reactome, KEGG, and Panther pathway resources, focusing on a curated set of neuronal/synaptic signaling, neurotransmitter cycle, axon guidance, oxidative stress/FOXO signaling, and neurodegeneration-related pathway themes (including Alzheimer’s- and Parkinson’s-relevant pathway collections, Supplemental Table 2). RNA-seq outcomes were then pruned to match the resulting pathway- derived gene lists, and the retained genes were ranked by the opposition index cluster truncation at cluster 7; plus retaining MME and MAPT.

The selected Alzheimer’s-directed gene set was used to generate volcano plots summarizing differential behavior and a net treatment effect vs total magnitude plot summarizing net endpoint versus total magnitude across miRNA-antagomir behavior. The selected genes were also used as seed genes in the NetworkAnalyst web utility to construct minimal protein–protein interaction (PPI) networks using the hippocampus and brain frontal cortex databases.

Networks were then manually adjusted to incorporate miR-181 as a node linked to each seed gene and reincorporate any seed genes excluded during initial network construction. All genes present in each final network were used to query the KEGG database for pathway memberships. To reduce within-sample redundancy, pathway results were pruned using the rule: If all genes within one KEGG pathway hit were entirely contained within another KEGG pathway hit, the smaller (contained) pathway was dropped. This pruning was performed solely to avoid redundant reporting and is not to be considered a biological claim.

We then compared pathways by Jaccard distance computed from in-sample gene membership sets and clustered using partitioning around medoids, with cluster count selection evaluated using ASW criteria. Cluster robustness was assessed by bootstrap subsampling of the input gene list (drop-only perturbation), restricting observed pathway hit sets by intersection with each bootstrap subsample. Cluster-level distance matrices (mean and median between-cluster distances) were derived from the Jaccard distance matrix and cluster assignments, and enrichment “bubble” plots were generated at the cluster level to visualize enrichment strength and hit counts.

### Data and Statistical Analysis

Western blots were scanned, and target band densitometry was determined by ImageJ software. Analysis of variance (ANOVA) of generalized linear models (glm) was applied followed by estimated marginal means with false discovery rate adjustment for most data. Associatin of miR-181a-d and covariates with human subject diagnosis status was tested by ordinal logistic regression (olr) models. The olr regresses ordinal response data (in this case NCI/MCI/AD) vs. predictors, using log-odds derived from ordered combinations of proportions of each outcome.(*50*) Exponentiated coefficients can be used to predict probabilities of each possible outcome in the data vs. predictor variables. Significance for all analyses was set at p ≤ 0.05. Effect sizes for glm/ANOVA were calculated as ω^2^ (omega square) (*51*). Effect sizes for ols were calculated as a modified Tjur’s *D* (*52*) for logistic models. Analyses were performed using R and the cluster, clusterProfiler, cowplot, fpc, ggplot2, glmGamPoi, gtable, grid, igraph, org.Hs.eg.db, and shadowtext packages.

## Acknowledgments

Sincere thanks to Drs. Martin Farlow, Bernardino Ghetti, Justin Long, Nipun Chopra, and Andrew Saykin.

This work was supported by the National Institute on Aging (NIA), National Institutes of Health (NIH), under award numbers P01-AG014449, R56-AG072810, R56-AG051086, R21-AG056007, R21-AG074539, and R21-AG076202.

We acknowledge the Research Education Core (REC) of Indiana Alzheimer’s Disease Research Center (NIA-P30 AG072976). The RNA-seq work performed in this work by the Indiana University School of Medicine Proteomics Core, and supported, in part, by the Indiana CTSI fund and by NIH award.

**We gratefully acknowledge that association studies of SNPs near the *MIR181A1* and *MIR181B1* and neprilysin genes with entorhinal cortex thickness were performed in ADNI cohort using the** ADNI database (adni.loni.usc.edu). ADNI was launched in 2003 as a public-private partnership led by Principal Investigator Michael W. Weiner, MD. The primary goal of ADNI has been to test whether serial MRI, PET, other biological markers, and clinical and neuropsychological assessment can be combined to measure the progression of mild cognitive impairment and early Alzheimer’s disease. Some Data collection and sharing for this project was funded by the ADNI (National Institutes of Health Grant U01 AG024904) and DOD ADNI (Department of Defense award number W81XWH-12-2-0012). ADNI is funded by the National Institute on Aging, the National Institute of Biomedical Imaging and Bioengineering, and through generous contributions from various private funders. We provide the full acknowledgement list (Supplemental Table 3).

## References

1. N. Iwata, S. Tsubuki, Y. Takaki, K. Watanabe, M. Sekiguchi, E. Hosoki, M. Kawashima-Morishima, H. J. Lee, E. Hama, Y. Sekine-Aizawa, T. C. Saido, Identification of the major Abeta1-42-degrading catabolic pathway in brain parenchyma: suppression leads to biochemical and pathological deposition. Nat Med 6, 143–150 (2000).

2. J. Hu, A. Igarashi, M. Kamata, H. Nakagawa, Angiotensin-converting enzyme degrades Alzheimer amyloid beta-peptide (A beta); retards A beta aggregation, deposition, fibril formation; and inhibits cytotoxicity. J Biol Chem 276, 47863–47868 (2001).

3. E. A. Eckman, M. Watson, L. Marlow, K. Sambamurti, C. B. Eckman, Alzheimer’s disease beta-amyloid peptide is increased in mice deficient in endothelin-converting enzyme. J Biol Chem 278, 2081–2084 (2003).

4. N. Carty, K. R. Nash, M. Brownlow, D. Cruite, D. Wilcock, M. L. Selenica, D. C. Lee, M. N. Gordon, D. Morgan, Intracranial injection of AAV expressing NEP but not IDE reduces amyloid pathology in APP+PS1 transgenic mice. PLoS One 8, e59626 (2013).

5. N. Iwata, S. Tsubuki, Y. Takaki, K. Shirotani, B. Lu, N. P. Gerard, C. Gerard, E. Hama, H. J. Lee, T. C. Saido, Metabolic regulation of brain Abeta by neprilysin. Science 292, 1550–1552 (2001).

6. S. L. Ameres, P. D. Zamore, Diversifying microRNA sequence and function. Nat Rev Mol Cell Biol 14, 475–488 (2013).

7. J. Krol, I. Loedige, W. Filipowicz, The widespread regulation of microRNA biogenesis, function and decay. Nat Rev Genet 11, 597–610 (2010).

8. M. R. Fabian, N. Sonenberg, W. Filipowicz, Regulation of mRNA translation and stability by microRNAs. Annu Rev Biochem 79, 351–379 (2010).

9. A. Bell-Hensley, S. Das, A. McAlinden, The miR-181 family: Wide-ranging pathophysiological effects on cell fate and function. J Cell Physiol 238, 698–713 (2023).

10. A. Ansari, E. Maffioletti, E. Milanesi, M. Marizzoni, G. B. Frisoni, O. Blin, J. C. Richardson, R. Bordet, G. Forloni, M. Gennarelli, L. Bocchio-Chiavetto, miR-146a and miR-181a are involved in the progression of mild cognitive impairment to Alzheimer’s disease. Neurobiol Aging 82, 102–109 (2019).

11. J. P. Cogswell, J. Ward, I. A. Taylor, M. Waters, Y. Shi, B. Cannon, K. Kelnar, J. Kemppainen, D. Brown, C. Chen, R. K. Prinjha, J. C. Richardson, A. M. Saunders, A. D. Roses, C. A. Richards, Identification of miRNA changes in Alzheimer’s disease brain and CSF yields putative biomarkers and insights into disease pathways. J Alzheimers Dis 14, 27–41 (2008).

12. P. Takousis, A. Sadlon, J. Schulz, I. Wohlers, V. Dobricic, L. Middleton, C. M. Lill, R. Perneczky, L. Bertram, Differential expression of microRNAs in Alzheimer’s disease brain, blood, and cerebrospinal fluid. Alzheimers Dement 15, 1468–1477 (2019).

13. C. J. Rodriguez-Ortiz, D. Baglietto-Vargas, H. Martinez-Coria, F. M. LaFerla, M. Kitazawa, Upregulation of miR-181 decreases c-Fos and SIRT-1 in the hippocampus of 3xTg-AD mice. J Alzheimers Dis 42, 1229–1238 (2014).

14. H. Geekiyanage, C. Chan, MicroRNA-137/181c regulates serine palmitoyltransferase and in turn amyloid β, novel targets in sporadic Alzheimer’s disease. J Neurosci 31, 14820–14830 (2011).

15. J. A. Bailey, B. Maloney, Y. W. Ge, D. K. Lahiri, Functional activity of the novel Alzheimer’s amyloid β-peptide interacting domain (AβID) in the APP and BACE1 promoter sequences and implications in activating apoptotic genes and in amyloidogenesis. Gene 488, 13–22 (2011).

16. B. Maloney, D. K. Lahiri, The Alzheimer’s amyloid β-peptide (Aβ) binds a specific DNA Aβ-interacting domain (AβID) in the APP, BACE1, and APOE promoters in a sequence-specific manner: characterizing a new regulatory motif. Gene 488, 1–12 (2011).

17. A. Indrieri, S. Carrella, P. Carotenuto, S. Banfi, B. Franco, The Pervasive Role of the miR-181 Family in Development, Neurodegeneration, and Cancer. Int J Mol Sci 21, (2020).

18. L. Lebson, K. Nash, S. Kamath, D. Herber, N. Carty, D. C. Lee, Q. Li, K. Szekeres, U. Jinwal, J. Koren, C. A. Dickey, P. E. Gottschall, D. Morgan, M. N. Gordon, Trafficking CD11b-positive blood cells deliver therapeutic genes to the brain of amyloid-depositing transgenic mice. J Neurosci 30, 9651–9658 (2010).

19. Y. Li, Y. Wang, J. Wang, K. Y. Chong, J. Xu, Z. Liu, C. Shan, Expression of Neprilysin in Skeletal Muscle by Ultrasound-Mediated Gene Transfer (Sonoporation) Reduces Amyloid Burden for AD. Mol Ther Methods Clin Dev 17, 300–308 (2020).

20. J. Avila, J. J. Lucas, M. Perez, F. Hernandez, Role of tau protein in both physiological and pathological conditions. Physiol Rev 84, 361–384 (2004).

21. A. Caceres, K. S. Kosik, Inhibition of neurite polarity by tau antisense oligonucleotides in primary cerebellar neurons. Nature 343, 461–463 (1990).

22. M. Mundalil Vasu, A. Anitha, I. Thanseem, K. Suzuki, K. Yamada, T. Takahashi, T. Wakuda, K. Iwata, M. Tsujii, T. Sugiyama, N. Mori, Serum microRNA profiles in children with autism. Mol Autism 5, 40 (2014).

23. C. M. Schumann, F. R. Sharp, B. P. Ander, B. Stamova, Possible sexually dimorphic role of miRNA and other sncRNA in ASD brain. Mol Autism 8, 4 (2017).

24. A. Kos, N. Olde Loohuis, J. Meinhardt, H. van Bokhoven, B. B. Kaplan, G. J. Martens, A. Aschrafi, MicroRNA-181 promotes synaptogenesis and attenuates axonal outgrowth in cortical neurons. Cell Mol Life Sci 73, 3555–3567 (2016).

25. C. Braicu, F. D. Mureșanu, E. Isachesku, N. Bornstein, S. R. Filipović, S. Strilciuc, A. Pana, Role of miR-181 Family Members in Stroke: Insights into Mechanisms and Therapeutic Potential. Int J Mol Sci 26, (2025).

26. C. S. Stein, J. M. McLendon, N. H. Witmer, R. L. Boudreau, Modulation of miR-181 influences dopaminergic neuronal degeneration in a mouse model of Parkinson’s disease. Mol Ther Nucleic Acids 28, 1–15 (2022).

27. F. Noorbakhsh, K. K. Ellestad, F. Maingat, K. G. Warren, M. H. Han, L. Steinman, G. B. Baker, C. Power, Impaired neurosteroid synthesis in multiple sclerosis. Brain 134, 2703–2721 (2011).

28. L. J. Xu, Y. B. Ouyang, X. Xiong, C. M. Stary, R. G. Giffard, Post-stroke treatment with miR-181 antagomir reduces injury and improves long-term behavioral recovery in mice after focal cerebral ischemia. Exp Neurol 264, 1–7 (2015).

29. I. Magen, N. S. Yacovzada, E. Yanowski, A. Coenen-Stass, J. Grosskreutz, C. H. Lu, L. Greensmith, A. Malaspina, P. Fratta, E. Hornstein, Circulating miR-181 is a prognostic biomarker for amyotrophic lateral sclerosis. Nat Neurosci 24, 1534–1541 (2021).

30. Y. Cohen, I. Sinai, I. Magen, Y. M. Danino, J. Wuu, A. Malaspina, M. Benatar, E. Hornstein, IsomiR utility in amyotrophic lateral sclerosis prognostication. Med 7, 100928 (2026).

31. T. W. Mills, M. Wu, J. Alonso, H. Puente, J. Charles, Z. Chen, S. H. Yoo, M. D. Mayes, S. Assassi, Unraveling the role of MiR-181 in skin fibrosis pathogenesis by targeting NUDT21. Faseb j 38, e70022 (2024).

32. J. Chen, K. Liu, M. A. Vadas, J. R. Gamble, G. W. McCaughan, The Role of the MiR-181 Family in Hepatocellular Carcinoma. Cells 13, (2024).

33. G. Singh, S. Singh, I. Mohapatra, S. Kim, M. Sharma, J. Akers, T. Nguyen, E. Wong, M. M. Moreno, E. Kokkoli, S. Vasudevan, S. E. Lawler, W. S. El-Deiry, Z. Gokaslan, C. C. Chen, Feedforward miR-181d degradation modulates population variance of methyl-guanine methyl transferase and temozolomide resistance. Cell Rep 44, 116516 (2025).

34. G. Singh, S. Singh, I. Mohapatra, J. Hou, A. Ni, D. Barik, H. Zheng, S. Kim, M. Sharma, S. Lawler, S. Vasudevan, E. Kokkoli, S. Sarangi, H. Elinzano, E. T. Wong, M. Martinez-Moreno, Z. Gokaslan, W. El-Deiry, C. C. Chen, miR-181d coordinates homologous recombination and anti-tumor immune responses in glioblastoma. iScience 29, 115077 (2026).

35. Y. B. Ouyang, Y. Lu, S. Yue, R. G. Giffard, miR-181 targets multiple Bcl-2 family members and influences apoptosis and mitochondrial function in astrocytes. Mitochondrion 12, 213–219 (2012).

36. K. A. Tekirdag, G. Korkmaz, D. G. Ozturk, R. Agami, D. Gozuacik, MIR181A regulates starvation- and rapamycin-induced autophagy through targeting of ATG5. Autophagy 9, 374–385 (2013).

37. J. Guo, Y. Ma, X. Peng, H. Jin, J. Liu, LncRNA CCAT1 promotes autophagy via regulating ATG7 by sponging miR-181 in hepatocellular carcinoma. J Cell Biochem 120, 17975–17983 (2019).

38. E. R. Hutchison, E. M. Kawamoto, D. D. Taub, A. Lal, K. Abdelmohsen, Y. Zhang, W. H. Wood, 3rd, E. Lehrmann, S. Camandola, K. G. Becker, M. Gorospe, M. P. Mattson, Evidence for miR-181 involvement in neuroinflammatory responses of astrocytes. Glia 61, 1018–1028 (2013).

39. C. M. Cahill, S. S. Sarang, R. Bakshi, N. Xia, D. K. Lahiri, J. T. Rogers, Neuroprotective Strategies and Cell-Based Biomarkers for Manganese-Induced Toxicity in Human Neuroblastoma (SH-SY5Y) Cells. Biomolecules 14, (2024).

40. A. Bermejo-Santos, M. Rubio-García, R. Torrillas-de la Cal, R. Casado-Navarro, E. Serrano-Saiz, Dmrt2 regulates sex-biased neuronal development in the cingulate cortex. Cell Mol Life Sci 82, 376 (2025).

41. V. Agarwal, G. W. Bell, J. W. Nam, D. P. Bartel, Predicting effective microRNA target sites in mammalian mRNAs. Elife 4, (2015).

42. M. D. Paraskevopoulou, G. Georgakilas, N. Kostoulas, I. S. Vlachos, T. Vergoulis, M. Reczko, C. Filippidis, T. Dalamagas, A. G. Hatzigeorgiou, DIANA-microT web server v5.0: service integration into miRNA functional analysis workflows. Nucleic Acids Res 41, W169–173 (2013).

43. M. Reczko, M. Maragkakis, P. Alexiou, I. Grosse, A. G. Hatzigeorgiou, Functional microRNA targets in protein coding sequences. Bioinformatics 28, 771–776 (2012).

44. W. Liu, X. Wang, Prediction of functional microRNA targets by integrative modeling of microRNA binding and target expression data. Genome Biol 20, 18 (2019).

45. Y. Chen, X. Wang, miRDB: an online database for prediction of functional microRNA targets. Nucleic Acids Res 48, D127–d131 (2020).

46. K. C. Miranda, T. Huynh, Y. Tay, Y. S. Ang, W. L. Tam, A. M. Thomson, B. Lim, I. Rigoutsos, A pattern-based method for the identification of MicroRNA binding sites and their corresponding heteroduplexes. Cell 126, 1203–1217 (2006).

47. C. E. Vejnar, E. M. Zdobnov, MiRmap: comprehensive prediction of microRNA target repression strength. Nucleic Acids Res 40, 11673–11683 (2012).

48. S. Kanoria, W. Rennie, C. Liu, C. S. Carmack, J. Lu, Y. Ding, STarMir: tools for prediction of microRNA binding sites. Methods Mol. Biol. 1490, 73–82 (2016).

49. R. Wang, B. Maloney, J. S. Beck, S. E. Counts, D. K. Lahiri, MicroRNA-153-3p targets repressor element 1-silencing transcription factor (REST) and neuronal differentiation: Implications for Alzheimer’s disease. Alzheimers Dement 21, e70399 (2025).

50. A. Agresti, “An Introduction to Categorical Data Analysis, 3rd Edition” (Wiley, Hoboken, NJ, USA, 2018), chap. Multicategory logit models, pp. 159–192.

51. S. Olejnik, J. Algina, Generalized eta and omega squared statistics: measures of effect size for some common research designs. Psychol Methods 8, 434–447 (2003).

52. T. Tjur, Coefficients of Determination in Logistic Regression Models—A New Proposal: The Coefficient of Discrimination. The American Statistician 63, 366–372 (2009).

53. P. H. Kuhn, H. Wang, B. Dislich, A. Colombo, U. Zeitschel, J. W. Ellwart, E. Kremmer, S. Rossner, S. F. Lichtenthaler, ADAM10 is the physiologically relevant, constitutive alpha-secretase of the amyloid precursor protein in primary neurons. Embo j 29, 3020–3032 (2010).

54. C. Haass, M. G. Schlossmacher, A. Y. Hung, C. Vigo-Pelfrey, A. Mellon, B. L. Ostaszewski, I. Lieberburg, E. H. Koo, D. Schenk, D. B. Teplow, et al., Amyloid beta- peptide is produced by cultured cells during normal metabolism. Nature 359, 322–325 (1992).

55. R. Yan, J. B. Munzner, M. E. Shuck, M. J. Bienkowski, BACE2 functions as an alternative alpha-secretase in cells. J Biol Chem 276, 34019–34027 (2001).

56. Y. Mimori-Kiyosue, I. Grigoriev, G. Lansbergen, H. Sasaki, C. Matsui, F. Severin, N. Galjart, F. Grosveld, I. Vorobjev, S. Tsukita, A. Akhmanova, CLASP1 and CLASP2 bind to EB1 and regulate microtubule plus-end dynamics at the cell cortex. J Cell Biol 168, 141–153 (2005).

57. M. Furukawa, Y. J. He, C. Borchers, Y. Xiong, Targeting of protein ubiquitination by BTB-Cullin 3-Roc1 ubiquitin ligases. Nat. Cell Biol. 5, 1001–1007 (2003).

58. T. Wahle, D. R. Thal, M. Sastre, A. Rentmeister, N. Bogdanovic, M. Famulok, M. T. Heneka, J. Walter, GGA1 is expressed in the human brain and affects the generation of amyloid beta-peptide. J Neurosci 26, 12838–12846 (2006).

59. M. Shibata, S. Yamada, S. R. Kumar, M. Calero, J. Bading, B. Frangione, D. M. Holtzman, C. A. Miller, D. K. Strickland, J. Ghiso, B. V. Zlokovic, Clearance of Alzheimer’s amyloid-ss(1-40) peptide from brain by LDL receptor-related protein-1 at the blood-brain barrier. J Clin Invest 106, 1489–1499 (2000).

60. T. Ohta, J. J. Michel, A. J. Schottelius, Y. Xiong, ROC1, a homolog of APC11, represents a family of cullin partners with an associated ubiquitin ligase activity. Mol Cell 3, 535–541 (1999).

61. A. W. Schmid, E. Condemi, G. Tuchscherer, D. Chiappe, M. Mutter, H. Vogel, M. Moniatte, Y. O. Tsybin, Tissue transglutaminase-mediated glutamine deamidation of beta-amyloid peptide increases peptide solubility, whereas enzymatic cross-linking and peptide fragmentation may serve as molecular triggers for rapid peptide aggregation. J Biol Chem 286, 12172–12188 (2011).

62. M. Lu, T. Liu, Q. Jiao, J. Ji, M. Tao, Y. Liu, Q. You, Z. Jiang, Discovery of a Keap1-dependent peptide PROTAC to knockdown Tau by ubiquitination-proteasome degradation pathway. Eur. J. Med. Chem. 146, 251–259 (2018).

63. C. A. Ferré, A. Thouard, A. Bétourné, A. L. Le Dorze, P. Belenguer, M. C. Miquel, J. M. Peyrin, D. Gonzalez-Dunia, M. Szelechowski, HSPA9/Mortalin mediates axo-protection and modulates mitochondrial dynamics in neurons. Sci Rep 11, 17705 (2021).

64. K. S. Kosik, C. L. Joachim, D. J. Selkoe, Microtubule-associated protein tau (tau) is a major antigenic component of paired helical filaments in Alzheimer disease. Proc Natl Acad Sci U S A 83, 4044–4048 (1986).

65. J. Biernat, Y. Z. Wu, T. Timm, Q. Zheng-Fischhöfer, E. Mandelkow, L. Meijer, E. M. Mandelkow, Protein kinase MARK/PAR-1 is required for neurite outgrowth and establishment of neuronal polarity. Mol Biol Cell 13, 4013–4028 (2002).

66. K. Nakamura, A. Greenwood, L. Binder, E. H. Bigio, S. Denial, L. Nicholson, X. Z. Zhou, K. P. Lu, Proline isomer-specific antibodies reveal the early pathogenic tau conformation in Alzheimer’s disease. Cell 149, 232–244 (2012).

67. S. Luo, H. Liu, T. Xiao, Y. Li, X. Liu, X. Xiao, X. Liao, Y. Liu, Y. Zhou, J. L. Wang, J. Guo, T. Tu, X. Yan, B. Tang, Z. Zhang, B. Jiao, L. Shen, Neuronal PPP2R5C in plasma is a potential biomarker for early diagnosis of Alzheimer’s disease. Cell Rep Med 7, 102631 (2026).

68. H. B. Luo, Y. Y. Xia, X. J. Shu, Z. C. Liu, Y. Feng, X. H. Liu, G. Yu, G. Yin, Y. S. Xiong, K. Zeng, J. Jiang, K. Ye, X. C. Wang, J. Z. Wang, SUMOylation at K340 inhibits tau degradation through deregulating its phosphorylation and ubiquitination. Proc Natl Acad Sci U S A 111, 16586–16591 (2014).

69. R. A. Halverson, J. Lewis, S. Frausto, M. Hutton, N. A. Muma, Tau protein is cross-linked by transglutaminase in P301L tau transgenic mice. J Neurosci 25, 1226–1233 (2005).

70. N. F. Darwich, J. M. Phan, B. Kim, E. Suh, J. D. Papatriantafyllou, L. Changolkar, A. T. Nguyen, C. M. O’Rourke, Z. He, S. Porta, G. S. Gibbons, K. C. Luk, S. G. Papageorgiou, M. Grossman, L. Massimo, D. J. Irwin, C. T. McMillan, I. M. Nasrallah, C. Toro, G. K. Aguirre, V. M. Van Deerlin, E. B. Lee, Autosomal dominant VCP hypomorph mutation impairs disaggregation of PHF-tau. Science 370, (2020).

71. H. Weidberg, E. Shvets, T. Shpilka, F. Shimron, V. Shinder, Z. Elazar, LC3 and GATE-16/GABARAP subfamilies are both essential yet act differently in autophagosome biogenesis. Embo j 29, 1792–1802 (2010).

72. E. J. Bae, D. K. Kim, C. Kim, M. Mante, A. Adame, E. Rockenstein, A. Ulusoy, M. Klinkenberg, G. R. Jeong, J. R. Bae, C. Lee, H. J. Lee, B. D. Lee, D. A. Di Monte, E. Masliah, S. J. Lee, LRRK2 kinase regulates α-synuclein propagation via RAB35 phosphorylation. Nat Commun 9, 3465 (2018).

73. C. Bussi, J. M. Peralta Ramos, D. S. Arroyo, J. I. Gallea, P. Ronchi, A. Kolovou, J. M. Wang, O. Florey, M. S. Celej, Y. Schwab, N. T. Ktistakis, P. Iribarren, Alpha-synuclein fibrils recruit TBK1 and OPTN to lysosomal damage sites and induce autophagy in microglial cells. J Cell Sci 131, (2018).

74. H. Shimura, M. G. Schlossmacher, N. Hattori, M. P. Frosch, A. Trockenbacher, R. Schneider, Y. Mizuno, K. S. Kosik, D. J. Selkoe, Ubiquitination of a new form of alpha- synuclein by parkin from human brain: implications for Parkinson’s disease. Science 293, 263–269 (2001).

75. M. G. Spillantini, M. L. Schmidt, V. M. Lee, J. Q. Trojanowski, R. Jakes, M. Goedert, Alpha-synuclein in Lewy bodies. Nature 388, 839–840 (1997).

76. Y. Watanabe, H. Tatebe, K. Taguchi, Y. Endo, T. Tokuda, T. Mizuno, M. Nakagawa, M. Tanaka, p62/SQSTM1-dependent autophagy of Lewy body-like α-synuclein inclusions. PLoS One 7, e52868 (2012).

77. R. Rott, R. Szargel, V. Shani, H. Hamza, M. Savyon, F. Abd Elghani, R. Bandopadhyay, S. Engelender, SUMOylation and ubiquitination reciprocally regulate α-synuclein degradation and pathological aggregation. Proc Natl Acad Sci U S A 114, 13176–13181 (2017).

78. R. Suzuki, H. Kawahara, UBQLN4 recognizes mislocalized transmembrane domain proteins and targets these to proteasomal degradation. EMBO Rep 17, 842–857 (2016).

79. J. Zhu, S. Pittman, D. Dhavale, R. French, J. N. Patterson, M. S. Kaleelurrrahuman, Y. Sun, J. Vaquer-Alicea, G. Maggiore, C. S. Clemen, W. J. Buscher, J. Bieschke, P. Kotzbauer, Y. Ayala, M. I. Diamond, A. A. Davis, C. Weihl, VCP suppresses proteopathic seeding in neurons. Mol Neurodegener 17, 30 (2022).

