## Supplemental Materials for "MicroRNA-181 influences Alzheimer’s risk by regulating neprilysin and microtubule-associated tau pathways, offering a novel target"

### Supplemental Material

#### Supplemental Table 1. Genes directly involved in Aβ, p-tau, or α-synuclein aggregation or clearance.

| o | Pathology | Activity | Evidence type | Ref. |
|---|---|---|---|---|
| ADAM10 | Aβ | Physiological α-secretase cleavage of APP; non-amyloidogenic APP processing that prevents/reduces Aβ generation | Pathology-specific | (*53*) |
| APP | Aβ | APP is proteolytically processed to produce Aβ peptides | Pathology-specific | (*54*) |
| BACE2 | Aβ | APP-processing β-secretase homolog; in cells, BACE2 acts mainly as an alternative α-secretase-like APP-processing enzyme | Pathology/activity-specific; wording corrected | (*55*) |
| CLASP2 | Aβ | Microtubule plus-end tracking protein regulating microtubule plus-end dynamics | Activity-only | (*56*) |
| CUL3 | Aβ | CUL3 E3 ligase can ubiquitinate and promote degradation of tau when recruited via Keap1 | Activity-only | (*57*) |
| GGA1 | Aβ | Adaptor affecting APP/BACE1 trafficking and Aβ generation | Pathology-specific | (*58*) |
| LRP1 | Aβ | Receptor mediating BBB/brain clearance of Aβ | Pathology-specific | (*59*) |
| MME | Aβ | Neprilysin metabolically regulates/degrades brain Aβ | Pathology-specific | (*5*) |
| RBX1 | Aβ | Cullin RING partner with ubiquitin ligase activity | Activity-only | (*60*) |
| TGM2 | Aβ | Tissue transglutaminase covalently modifies/cross-links Aβ peptides and can influence aggregation | Pathology-specific | (*61*) |
| CUL3 | Tau | Engineered Keap1–CUL3 recruitment can promote tau ubiquitination/degradation; not an endogenous canonical CUL3–tau pathway | Engineered tau-specific | (*62*) |
| HSPA9 | Tau | Mortalin/HSPA9 is a mitochondrial Hsp70 chaperone involved in mitochondrial protein import/folding quality control and neuronal stress | Activity-only | (*63*) |
| MAPT | Tau | Tau is a major component of paired helical filaments/neurofibrillary tangles | Pathology-specific | (*64*) |
| MARK2 | Tau | MARK/PAR-1 family kinases phosphorylate tau at KXGS motifs in the repeat domain, reducing tau–microtubule affinity | Pathology/activity-specific | (*65*) |
| PIN1 | Tau | Prolyl isomerase converting pathogenic cis phosphorylated tau toward trans p-tau, limiting tau pathology | Pathology-specific | (*66*) |
| PPP2R5C | Tau | PPP2R5C interacts with tau and reduces tau phosphorylation and abundance via PP2A and ULK1-dependent autophagolysosomal regulation. | Pathology-specific | (*67*) |
| RBX1 | Tau | Cullin RING partner with ubiquitin ligase activity | Activity-only | (*60*) |
| SUMO1 | Tau | SUMO1-mediated tau SUMOylation at K340 inhibits tau degradation and alters phosphorylation/ubiquitination/solubility | Pathology-specific | (*68*) |
| TGM2 | Tau | Transglutaminase cross-links tau into stable/insoluble aggregates | Pathology-specific | (*69*) |
| VCP | Tau | AAA+ ATPase/disaggregase activity against PHF-tau; VCP hypomorph mutation impairs tau disaggregation | Pathology-specific | (*70*) |
| GABARAPL2 | α-Syn | ATG8/GABARAP-family autophagy protein required for later autophagosome maturation | Activity-only | (*71*) |
| LRRK2 | α-Syn | LRRK2 kinase regulates α-synuclein propagation through RAB35 phosphorylation and lysosomal/vesicular pathways | Pathology-specific | (*72*) |

| o | Pathology | Activity | Evidence type | Ref. |
|---|---|---|---|---|
| OPTN | α-Syn | α-synuclein fibrils induce lysosomal damage that recruits TBK1/OPTN to ubiquitylated lysosomes and triggers an autophagic/lysophagy response; not direct α-syn cargo targeting | Pathology-context; lysophagy-specific | (*73*) |
| PRKN | α-Syn | Parkin ubiquitinates a 22-kDa glycosylated α-synuclein species | Pathology-specific | (*74*) |
| SNCA | α-Syn | α-synuclein is a major component of Lewy bodies | Pathology-specific | (*75*) |
| SQSTM1 | α-Syn | p62/SQSTM1 surrounds ubiquitinated Lewy-body-like α-synuclein inclusions and directs them to the autophagy–lysosome pathway | Pathology-specific | (*76*) |
| SUMO1 | α-Syn | SUMOylation regulates α-synuclein degradation and pathological aggregation | Pathology-specific | (*77*) |
| UBQLN4 | α-Syn | UBQLN4 recognizes misassembled/mislocalized proteins and targets them to proteasomal degradation | Activity-only | (*78*) |
| VCP | α-Syn | VCP suppresses proteopathic α-synuclein seeding and spread. | Pathology-specific | (*79*) |

**Supplemental Table 2. Summaries and Cluster definitions for hippocampus and frontal cortex KEGG pathways.**

| Brain Region | Pathway | Total | Hits | Expected | Hits/Exp. | FDR | Cluster |
|---|---|---|---|---|---|---|---|
| Hippocampus | Pathways of neurodegeneration - multiple diseases | 480 | 25 | 1.280 | 4.29 | < 0.001 | Neurodegeneration & Proteostasis |
| Hippocampus | Alzheimer disease | 388 | 22 | 0.910 | 4.59 | < 0.001 | Neurodegeneration & Proteostasis |
| Hippocampus | Ubiquitin mediated proteolysis | 142 | 11 | 0.167 | 6.05 | < 0.001 | Neurodegeneration & Proteostasis |
| Hippocampus | Mitophagy - animal | 105 | 9 | 0.101 | 6.48 | < 0.001 | Neurodegeneration & Proteostasis |
| Hippocampus | Parkinson disease | 268 | 14 | 0.400 | 5.13 | < 0.001 | Neurodegeneration & Proteostasis |
| Hippocampus | Amyotrophic lateral sclerosis | 368 | 14 | 0.549 | 4.67 | 0.002 | Neurodegeneration & Proteostasis |
| Hippocampus | Huntington disease | 308 | 10 | 0.328 | 4.93 | 0.020 | Neurodegeneration & Proteostasis |
| Hippocampus | Neurotrophin signaling pathway | 120 | 13 | 0.166 | 6.29 | < 0.001 | Immune & Stress Signaling |
| Hippocampus | MAPK signaling pathway | 300 | 18 | 0.576 | 4.97 | < 0.001 | Immune & Stress Signaling |
| Hippocampus | Autophagy - animal | 169 | 12 | 0.216 | 5.79 | < 0.001 | Immune & Stress Signaling |
| Hippocampus | Fluid shear stress and atherosclerosis | 142 | 11 | 0.167 | 6.05 | < 0.001 | Immune & Stress Signaling |
| Hippocampus | PD-L1 expression and PD-1 checkpoint pathway in cancer | 90 | 8 | 0.077 | 6.70 | < 0.001 | Immune & Stress Signaling |
| Hippocampus | Sphingolipid signaling pathway | 125 | 9 | 0.120 | 6.23 | < 0.001 | Immune & Stress Signaling |
| Hippocampus | NF-kappa B signaling pathway | 105 | 8 | 0.090 | 6.48 | < 0.001 | Immune & Stress Signaling |
| Hippocampus | Apoptosis | 137 | 9 | 0.132 | 6.10 | < 0.001 | Immune & Stress Signaling |
| Hippocampus | Osteoclast differentiation | 142 | 9 | 0.136 | 6.05 | < 0.001 | Immune & Stress Signaling |
| Hippocampus | Alcoholic liver disease | 144 | 9 | 0.138 | 6.02 | < 0.001 | Immune & Stress Signaling |
| Hippocampus | Lipid and atherosclerosis | 216 | 11 | 0.253 | 5.44 | < 0.001 | Immune & Stress Signaling |
| Hippocampus | RIG-I-like receptor signaling pathway | 72 | 6 | 0.046 | 7.02 | 0.001 | Immune & Stress Signaling |
| Hippocampus | C-type lectin receptor signaling pathway | 105 | 7 | 0.078 | 6.48 | 0.002 | Immune & Stress Signaling |
| Hippocampus | Th17 cell differentiation | 109 | 7 | 0.081 | 6.43 | 0.002 | Immune & Stress Signaling |
| Hippocampus | TNF signaling pathway | 119 | 7 | 0.089 | 6.30 | 0.003 | Immune & Stress Signaling |
| Hippocampus | T cell receptor signaling pathway | 122 | 7 | 0.091 | 6.26 | 0.004 | Immune & Stress Signaling |
| Hippocampus | mTOR signaling pathway | 158 | 8 | 0.135 | 5.89 | 0.004 | Immune & Stress Signaling |
| Hippocampus | IL-17 signaling pathway | 94 | 6 | 0.060 | 6.64 | 0.004 | Immune & Stress Signaling |
| Hippocampus | IgSF CAM signaling | 299 | 11 | 0.351 | 4.97 | 0.007 | Immune & Stress Signaling |
| Hippocampus | Insulin resistance | 109 | 6 | 0.070 | 6.43 | 0.008 | Immune & Stress Signaling |
| Hippocampus | Chemical carcinogenesis - reactive oxygen species | 227 | 8 | 0.194 | 5.37 | 0.025 | Immune & Stress Signaling |
| Hippocampus | NOD-like receptor signaling pathway | 187 | 7 | 0.140 | 5.65 | 0.028 | Immune & Stress Signaling |
| Hippocampus | Prostate cancer | 106 | 16 | 0.181 | 6.47 | < 0.001 | Oncogenes & Cell Fate |
| Hippocampus | PI3K-Akt signaling pathway | 362 | 25 | 0.965 | 4.69 | < 0.001 | Oncogenes & Cell Fate |
| Hippocampus | Gastric cancer | 150 | 16 | 0.256 | 5.97 | < 0.001 | Oncogenes & Cell Fate |
| Hippocampus | Cell cycle | 158 | 16 | 0.270 | 5.89 | < 0.001 | Oncogenes & Cell Fate |
| Hippocampus | Acute myeloid leukemia | 68 | 11 | 0.080 | 7.11 | < 0.001 | Oncogenes & Cell Fate |
| Hippocampus | Chronic myeloid leukemia | 77 | 11 | 0.090 | 6.93 | < 0.001 | Oncogenes & Cell Fate |
| Hippocampus | Breast cancer | 148 | 14 | 0.221 | 5.99 | < 0.001 | Oncogenes & Cell Fate |
| Hippocampus | Small cell lung cancer | 93 | 11 | 0.109 | 6.66 | < 0.001 | Oncogenes & Cell Fate |
| Hippocampus | Hippo signaling pathway | 157 | 13 | 0.218 | 5.90 | < 0.001 | Oncogenes & Cell Fate |
| Hippocampus | Proteoglycans in cancer | 204 | 14 | 0.305 | 5.52 | < 0.001 | Oncogenes & Cell Fate |
| Hippocampus | Cellular senescence | 157 | 12 | 0.201 | 5.90 | < 0.001 | Oncogenes & Cell Fate |
| Hippocampus | HIF-1 signaling pathway | 110 | 10 | 0.117 | 6.41 | < 0.001 | Oncogenes & Cell Fate |
| Hippocampus | Renal cell carcinoma | 70 | 8 | 0.060 | 7.07 | < 0.001 | Oncogenes & Cell Fate |
| Hippocampus | Adherens junction | 93 | 9 | 0.089 | 6.66 | < 0.001 | Oncogenes & Cell Fate |
| Hippocampus | Non-small cell lung cancer | 73 | 8 | 0.062 | 7.00 | < 0.001 | Oncogenes & Cell Fate |
| Hippocampus | Pancreatic cancer | 77 | 8 | 0.066 | 6.93 | < 0.001 | Oncogenes & Cell Fate |
| Hippocampus | Cadherin signaling | 404 | 18 | 0.776 | 4.54 | < 0.001 | Oncogenes & Cell Fate |
| Hippocampus | FoxO signaling pathway | 133 | 10 | 0.142 | 6.14 | < 0.001 | Oncogenes & Cell Fate |
| Hippocampus | Longevity regulating pathway | 90 | 8 | 0.077 | 6.70 | < 0.001 | Oncogenes & Cell Fate |
| Hippocampus | MicroRNAs in cancer | 323 | 15 | 0.517 | 4.86 | < 0.001 | Oncogenes & Cell Fate |
| Hippocampus | Cushing syndrome | 155 | 10 | 0.165 | 5.92 | < 0.001 | Oncogenes & Cell Fate |
| Hippocampus | Wnt signaling pathway | 178 | 10 | 0.190 | 5.72 | < 0.001 | Oncogenes & Cell Fate |
| Hippocampus | Axon guidance | 184 | 10 | 0.196 | 5.67 | < 0.001 | Oncogenes & Cell Fate |
| Hippocampus | Thyroid hormone signaling pathway | 122 | 8 | 0.104 | 6.26 | < 0.001 | Oncogenes & Cell Fate |
| Hippocampus | TGF-beta signaling pathway | 110 | 7 | 0.082 | 6.41 | 0.002 | Oncogenes & Cell Fate |

| Brain Region | Pathway | Total | Hits | Expected | Hits/Exp. | FDR | Cluster |
|---|---|---|---|---|---|---|---|
| Hippocampus | Hedgehog signaling pathway | 56 | 5 | 0.030 | 7.39 | 0.003 | Oncogenes & Cell Fate |
| Hippocampus | Transcriptional misregulation in cancer | 201 | 9 | 0.193 | 5.54 | 0.005 | Oncogenes & Cell Fate |
| Hippocampus | Focal adhesion | 203 | 9 | 0.195 | 5.53 | 0.005 | Oncogenes & Cell Fate |
| Hippocampus | JAK-STAT signaling pathway | 168 | 8 | 0.143 | 5.80 | 0.005 | Oncogenes & Cell Fate |
| Hippocampus | p53 signaling pathway | 75 | 5 | 0.040 | 6.97 | 0.008 | Oncogenes & Cell Fate |
| Hippocampus | Signaling pathways regulating pluripotency of stem cells | 144 | 7 | 0.108 | 6.02 | 0.008 | Oncogenes & Cell Fate |
| Hippocampus | Integrin signaling | 154 | 7 | 0.115 | 5.93 | 0.011 | Oncogenes & Cell Fate |
| Hippocampus | AMPK signaling pathway | 122 | 6 | 0.078 | 6.26 | 0.014 | Oncogenes & Cell Fate |
| Hippocampus | Oocyte meiosis | 138 | 13 | 0.191 | 6.09 | < 0.001 | Endocrine & Signal Transduction |
| Hippocampus | Endocrine resistance | 99 | 11 | 0.116 | 6.57 | < 0.001 | Endocrine & Signal Transduction |
| Hippocampus | Chemical carcinogenesis - receptor activation | 217 | 15 | 0.347 | 5.43 | < 0.001 | Endocrine & Signal Transduction |
| Hippocampus | Phospholipase D signaling pathway | 149 | 12 | 0.191 | 5.98 | < 0.001 | Endocrine & Signal Transduction |
| Hippocampus | GnRH signaling pathway | 93 | 9 | 0.089 | 6.66 | < 0.001 | Endocrine & Signal Transduction |
| Hippocampus | Glioma | 76 | 8 | 0.065 | 6.95 | < 0.001 | Endocrine & Signal Transduction |
| Hippocampus | Melanogenesis | 101 | 9 | 0.097 | 6.54 | < 0.001 | Endocrine & Signal Transduction |
| Hippocampus | cAMP signaling pathway | 226 | 12 | 0.289 | 5.37 | < 0.001 | Endocrine & Signal Transduction |
| Hippocampus | Estrogen signaling pathway | 139 | 9 | 0.133 | 6.08 | < 0.001 | Endocrine & Signal Transduction |
| Hippocampus | Rap1 signaling pathway | 212 | 11 | 0.249 | 5.47 | < 0.001 | Endocrine & Signal Transduction |
| Hippocampus | Longevity regulating pathway - multiple species | 62 | 6 | 0.040 | 7.24 | < 0.001 | Endocrine & Signal Transduction |
| Hippocampus | Gap junction | 89 | 7 | 0.066 | 6.72 | < 0.001 | Endocrine & Signal Transduction |
| Hippocampus | Growth hormone synthesis, secretion and action | 122 | 8 | 0.104 | 6.26 | < 0.001 | Endocrine & Signal Transduction |
| Hippocampus | AGE-RAGE signaling pathway in diabetic complications | 101 | 7 | 0.075 | 6.54 | 0.001 | Endocrine & Signal Transduction |
| Hippocampus | Parathyroid hormone synthesis, secretion and action | 115 | 7 | 0.086 | 6.35 | 0.003 | Endocrine & Signal Transduction |
| Hippocampus | Regulation of lipolysis in adipocytes | 59 | 5 | 0.031 | 7.31 | 0.003 | Endocrine & Signal Transduction |
| Hippocampus | Adrenergic signaling in cardiomyocytes | 154 | 8 | 0.131 | 5.93 | 0.003 | Endocrine & Signal Transduction |
| Hippocampus | Oxytocin signaling pathway | 154 | 8 | 0.131 | 5.93 | 0.003 | Endocrine & Signal Transduction |
| Hippocampus | VEGF signaling pathway | 60 | 5 | 0.032 | 7.29 | 0.003 | Endocrine & Signal Transduction |
| Hippocampus | GnRH secretion | 65 | 5 | 0.035 | 7.17 | 0.005 | Endocrine & Signal Transduction |
| Hippocampus | Calcium signaling pathway | 254 | 10 | 0.271 | 5.21 | 0.006 | Endocrine & Signal Transduction |
| Hippocampus | Insulin signaling pathway | 138 | 7 | 0.103 | 6.09 | 0.007 | Endocrine & Signal Transduction |
| Hippocampus | Vasopressin-regulated water reabsorption | 44 | 4 | 0.019 | 7.74 | 0.007 | Endocrine & Signal Transduction |
| Hippocampus | Platelet activation | 126 | 6 | 0.081 | 6.22 | 0.016 | Endocrine & Signal Transduction |
| Hippocampus | Inflammatory mediator regulation of TRP channels | 99 | 5 | 0.053 | 6.57 | 0.022 | Endocrine & Signal Transduction |
| Hippocampus | Endocytosis | 250 | 8 | 0.213 | 5.23 | 0.040 | Endocrine & Signal Transduction |
| Hippocampus | Relaxin signaling pathway | 130 | 13 | 0.180 | 6.17 | < 0.001 | Synaptic & Neuroactive Signaling |
| Hippocampus | Serotonergic synapse | 115 | 12 | 0.147 | 6.35 | < 0.001 | Synaptic & Neuroactive Signaling |
| Hippocampus | Circadian entrainment | 97 | 10 | 0.103 | 6.59 | < 0.001 | Synaptic & Neuroactive Signaling |

| Brain Region | Pathway | Total | Hits | Expected | Hits/Exp. | FDR | Cluster |
|---|---|---|---|---|---|---|---|
| Hippocampus | Ras signaling pathway | 238 | 15 | 0.381 | 5.30 | < 0.001 | Synaptic & Neuroactive Signaling |
| Hippocampus | Dopaminergic synapse | 133 | 11 | 0.156 | 6.14 | < 0.001 | Synaptic & Neuroactive Signaling |
| Hippocampus | Apelin signaling pathway | 140 | 11 | 0.164 | 6.07 | < 0.001 | Synaptic & Neuroactive Signaling |
| Hippocampus | Cholinergic synapse | 115 | 10 | 0.123 | 6.35 | < 0.001 | Synaptic & Neuroactive Signaling |
| Hippocampus | Glutamatergic synapse | 116 | 10 | 0.124 | 6.34 | < 0.001 | Synaptic & Neuroactive Signaling |
| Hippocampus | Chemokine signaling pathway | 193 | 12 | 0.247 | 5.60 | < 0.001 | Synaptic & Neuroactive Signaling |
| Hippocampus | Neuroactive ligand signaling | 199 | 12 | 0.255 | 5.56 | < 0.001 | Synaptic & Neuroactive Signaling |
| Hippocampus | GABAergic synapse | 89 | 8 | 0.076 | 6.72 | < 0.001 | Synaptic & Neuroactive Signaling |
| Hippocampus | Morphine addiction | 91 | 8 | 0.078 | 6.69 | < 0.001 | Synaptic & Neuroactive Signaling |
| Hippocampus | Alcoholism | 191 | 11 | 0.224 | 5.62 | < 0.001 | Synaptic & Neuroactive Signaling |
| Hippocampus | Retrograde endocannabinoid signaling | 149 | 9 | 0.143 | 5.98 | < 0.001 | Synaptic & Neuroactive Signaling |
| Hippocampus | Type II diabetes mellitus | 47 | 5 | 0.025 | 7.64 | 0.001 | Synaptic & Neuroactive Signaling |
| Hippocampus | Synaptic vesicle cycle | 79 | 6 | 0.051 | 6.89 | 0.002 | Synaptic & Neuroactive Signaling |
| Hippocampus | Hormone signaling | 219 | 10 | 0.234 | 5.42 | 0.002 | Synaptic & Neuroactive Signaling |
| Hippocampus | Cocaine addiction | 49 | 4 | 0.021 | 7.58 | 0.009 | Synaptic & Neuroactive Signaling |
| Hippocampus | Nicotine addiction | 41 | 3 | 0.013 | 7.84 | 0.033 | Synaptic & Neuroactive Signaling |
| Frontal Cortex | Alzheimer disease | 388 | 22 | 0.910 | 4.59 | < 0.001 | Neurodegeneration & Apoptosis |
| Frontal Cortex | Amyotrophic lateral sclerosis | 368 | 15 | 0.589 | 4.67 | < 0.001 | Neurodegeneration & Apoptosis |
| Frontal Cortex | Apoptosis | 137 | 11 | 0.161 | 6.10 | < 0.001 | Neurodegeneration & Apoptosis |
| Frontal Cortex | Fluid shear stress and atherosclerosis | 142 | 10 | 0.151 | 6.05 | < 0.001 | Neurodegeneration & Apoptosis |
| Frontal Cortex | Lipid and atherosclerosis | 216 | 14 | 0.323 | 5.44 | < 0.001 | Neurodegeneration & Apoptosis |
| Frontal Cortex | Longevity regulating pathway | 90 | 10 | 0.096 | 6.70 | < 0.001 | Neurodegeneration & Apoptosis |
| Frontal Cortex | MAPK signaling pathway | 300 | 20 | 0.640 | 4.97 | < 0.001 | Neurodegeneration & Apoptosis |
| Frontal Cortex | Mitophagy - animal | 105 | 9 | 0.101 | 6.48 | < 0.001 | Neurodegeneration & Apoptosis |
| Frontal Cortex | Neurotrophin signaling pathway | 120 | 14 | 0.179 | 6.29 | < 0.001 | Neurodegeneration & Apoptosis |
| Frontal Cortex | Pathways of neurodegeneration - multiple diseases | 480 | 25 | 1.280 | 4.29 | < 0.001 | Neurodegeneration & Apoptosis |
| Frontal Cortex | Small cell lung cancer | 93 | 11 | 0.109 | 6.66 | < 0.001 | Neurodegeneration & Apoptosis |
| Frontal Cortex | Sphingolipid signaling pathway | 125 | 9 | 0.120 | 6.23 | < 0.001 | Neurodegeneration & Apoptosis |
| Frontal Cortex | TNF signaling pathway | 119 | 8 | 0.102 | 6.30 | < 0.001 | Neurodegeneration & Apoptosis |
| Frontal Cortex | Adrenergic signaling in cardiomyocytes | 154 | 9 | 0.148 | 5.93 | < 0.001 | Synaptic Signaling |
| Frontal Cortex | Aldosterone synthesis and secretion | 98 | 5 | 0.052 | 6.58 | 0.023 | Synaptic Signaling |
| Frontal Cortex | Amphetamine addiction | 69 | 7 | 0.052 | 7.09 | < 0.001 | Synaptic Signaling |
| Frontal Cortex | cAMP signaling pathway | 226 | 12 | 0.289 | 5.37 | < 0.001 | Synaptic Signaling |

| Brain Region | Pathway | Total | Hits | Expected | Hits/Exp. | FDR | Cluster |
|---|---|---|---|---|---|---|---|
| Frontal Cortex | Chemical carcinogenesis - receptor activation | 217 | 16 | 0.370 | 5.43 | < 0.001 | Synaptic Signaling |
| Frontal Cortex | Endocrine resistance | 99 | 11 | 0.116 | 6.57 | < 0.001 | Synaptic Signaling |
| Frontal Cortex | Estrogen signaling pathway | 139 | 11 | 0.163 | 6.08 | < 0.001 | Synaptic Signaling |
| Frontal Cortex | Gap junction | 89 | 7 | 0.066 | 6.72 | < 0.001 | Synaptic Signaling |
| Frontal Cortex | Glucagon signaling pathway | 107 | 6 | 0.068 | 6.45 | 0.008 | Synaptic Signaling |
| Frontal Cortex | GnRH secretion | 65 | 5 | 0.035 | 7.17 | 0.005 | Synaptic Signaling |
| Frontal Cortex | GnRH signaling pathway | 93 | 10 | 0.099 | 6.66 | < 0.001 | Synaptic Signaling |
| Frontal Cortex | Growth hormone synthesis, secretion and action | 122 | 9 | 0.117 | 6.26 | < 0.001 | Synaptic Signaling |
| Frontal Cortex | Insulin secretion | 86 | 5 | 0.046 | 6.77 | 0.014 | Synaptic Signaling |
| Frontal Cortex | Insulin signaling pathway | 138 | 6 | 0.088 | 6.09 | 0.025 | Synaptic Signaling |
| Frontal Cortex | Longevity regulating pathway - multiple species | 62 | 8 | 0.053 | 7.24 | < 0.001 | Synaptic Signaling |
| Frontal Cortex | Long-term potentiation | 67 | 5 | 0.036 | 7.13 | 0.006 | Synaptic Signaling |
| Frontal Cortex | Melanogenesis | 101 | 9 | 0.097 | 6.54 | < 0.001 | Synaptic Signaling |
| Frontal Cortex | Oocyte meiosis | 138 | 13 | 0.191 | 6.09 | < 0.001 | Synaptic Signaling |
| Frontal Cortex | Oxytocin signaling pathway | 154 | 8 | 0.131 | 5.93 | 0.004 | Synaptic Signaling |
| Frontal Cortex | Parathyroid hormone synthesis, secretion and action | 115 | 8 | 0.098 | 6.35 | < 0.001 | Synaptic Signaling |
| Frontal Cortex | Parkinson disease | 268 | 14 | 0.400 | 5.13 | < 0.001 | Synaptic Signaling |
| Frontal Cortex | Phospholipase D signaling pathway | 149 | 13 | 0.207 | 5.98 | < 0.001 | Synaptic Signaling |
| Frontal Cortex | Protein processing in endoplasmic reticulum | 174 | 8 | 0.148 | 5.75 | 0.007 | Synaptic Signaling |
| Frontal Cortex | Rap1 signaling pathway | 212 | 10 | 0.226 | 5.47 | 0.002 | Synaptic Signaling |
| Frontal Cortex | Regulation of lipolysis in adipocytes | 59 | 5 | 0.031 | 7.31 | 0.003 | Synaptic Signaling |
| Frontal Cortex | Alcoholism | 191 | 13 | 0.265 | 5.62 | < 0.001 | Cancer & Cell Fate |
| Frontal Cortex | Apelin signaling pathway | 140 | 11 | 0.164 | 6.07 | < 0.001 | Cancer & Cell Fate |
| Frontal Cortex | Calcium signaling pathway | 254 | 10 | 0.271 | 5.21 | 0.007 | Cancer & Cell Fate |
| Frontal Cortex | Chemokine signaling pathway | 193 | 11 | 0.226 | 5.60 | < 0.001 | Cancer & Cell Fate |
| Frontal Cortex | Cholinergic synapse | 115 | 12 | 0.147 | 6.35 | < 0.001 | Cancer & Cell Fate |
| Frontal Cortex | Circadian entrainment | 97 | 10 | 0.103 | 6.59 | < 0.001 | Cancer & Cell Fate |
| Frontal Cortex | Cocaine addiction | 49 | 5 | 0.026 | 7.58 | 0.002 | Cancer & Cell Fate |
| Frontal Cortex | Dopaminergic synapse | 133 | 12 | 0.170 | 6.14 | < 0.001 | Cancer & Cell Fate |
| Frontal Cortex | Glutamatergic synapse | 116 | 10 | 0.124 | 6.34 | < 0.001 | Cancer & Cell Fate |
| Frontal Cortex | Hormone signaling | 219 | 10 | 0.234 | 5.42 | 0.003 | Cancer & Cell Fate |
| Frontal Cortex | Inflammatory mediator regulation of TRP channels | 99 | 5 | 0.053 | 6.57 | 0.024 | Cancer & Cell Fate |
| Frontal Cortex | Morphine addiction | 91 | 8 | 0.078 | 6.69 | < 0.001 | Cancer & Cell Fate |

| Brain Region | Pathway | Total | Hits | Expected | Hits/Exp. | FDR | Cluster |
|---|---|---|---|---|---|---|---|
| Frontal Cortex | Neuroactive ligand signaling | 199 | 12 | 0.255 | 5.56 | < 0.001 | Cancer & Cell Fate |
| Frontal Cortex | Nicotine addiction | 41 | 3 | 0.013 | 7.84 | 0.034 | Cancer & Cell Fate |
| Frontal Cortex | Ras signaling pathway | 238 | 15 | 0.381 | 5.30 | < 0.001 | Cancer & Cell Fate |
| Frontal Cortex | Relaxin signaling pathway | 130 | 13 | 0.180 | 6.17 | < 0.001 | Cancer & Cell Fate |
| Frontal Cortex | Retrograde endocannabinoid signaling | 149 | 9 | 0.143 | 5.98 | < 0.001 | Cancer & Cell Fate |
| Frontal Cortex | Serotonergic synapse | 115 | 12 | 0.147 | 6.35 | < 0.001 | Cancer & Cell Fate |
| Frontal Cortex | Synaptic vesicle cycle | 79 | 6 | 0.051 | 6.89 | 0.002 | Cancer & Cell Fate |
| Frontal Cortex | Vasopressin-regulated water reabsorption | 44 | 4 | 0.019 | 7.74 | 0.007 | Cancer & Cell Fate |
| Frontal Cortex | Acute myeloid leukemia | 68 | 10 | 0.073 | 7.11 | < 0.001 | Immune & Metabolic Stress |
| Frontal Cortex | Adherens junction | 93 | 10 | 0.099 | 6.66 | < 0.001 | Immune & Metabolic Stress |
| Frontal Cortex | Axon guidance | 184 | 10 | 0.196 | 5.67 | < 0.001 | Immune & Metabolic Stress |
| Frontal Cortex | Breast cancer | 148 | 14 | 0.221 | 5.99 | < 0.001 | Immune & Metabolic Stress |
| Frontal Cortex | Cadherin signaling | 404 | 18 | 0.776 | 4.54 | < 0.001 | Immune & Metabolic Stress |
| Frontal Cortex | Cell cycle | 158 | 15 | 0.253 | 5.89 | < 0.001 | Immune & Metabolic Stress |
| Frontal Cortex | Cellular senescence | 157 | 13 | 0.218 | 5.90 | < 0.001 | Immune & Metabolic Stress |
| Frontal Cortex | Chronic myeloid leukemia | 77 | 11 | 0.090 | 6.93 | < 0.001 | Immune & Metabolic Stress |
| Frontal Cortex | Cushing syndrome | 155 | 11 | 0.182 | 5.92 | < 0.001 | Immune & Metabolic Stress |
| Frontal Cortex | Focal adhesion | 203 | 10 | 0.217 | 5.53 | 0.002 | Immune & Metabolic Stress |
| Frontal Cortex | FoxO signaling pathway | 133 | 10 | 0.142 | 6.14 | < 0.001 | Immune & Metabolic Stress |
| Frontal Cortex | Gastric cancer | 150 | 16 | 0.256 | 5.97 | < 0.001 | Immune & Metabolic Stress |
| Frontal Cortex | Glioma | 76 | 8 | 0.065 | 6.95 | < 0.001 | Immune & Metabolic Stress |
| Frontal Cortex | Hedgehog signaling pathway | 56 | 5 | 0.030 | 7.39 | 0.003 | Immune & Metabolic Stress |
| Frontal Cortex | Hippo signaling pathway | 157 | 13 | 0.218 | 5.90 | < 0.001 | Immune & Metabolic Stress |
| Frontal Cortex | Integrin signaling | 154 | 7 | 0.115 | 5.93 | 0.012 | Immune & Metabolic Stress |
| Frontal Cortex | JAK-STAT signaling pathway | 168 | 8 | 0.143 | 5.80 | 0.006 | Immune & Metabolic Stress |
| Frontal Cortex | MicroRNAs in cancer | 323 | 16 | 0.551 | 4.86 | < 0.001 | Immune & Metabolic Stress |
| Frontal Cortex | Non-small cell lung cancer | 73 | 8 | 0.062 | 7.00 | < 0.001 | Immune & Metabolic Stress |
| Frontal Cortex | p53 signaling pathway | 75 | 5 | 0.040 | 6.97 | 0.008 | Immune & Metabolic Stress |
| Frontal Cortex | PI3K-Akt signaling pathway | 362 | 26 | 1.004 | 4.69 | < 0.001 | Immune & Metabolic Stress |
| Frontal Cortex | Prostate cancer | 106 | 17 | 0.192 | 6.47 | < 0.001 | Immune & Metabolic Stress |
| Frontal Cortex | Proteoglycans in cancer | 204 | 14 | 0.305 | 5.52 | < 0.001 | Immune & Metabolic Stress |
| Frontal Cortex | Renal cell carcinoma | 70 | 8 | 0.060 | 7.07 | < 0.001 | Immune & Metabolic Stress |
| Frontal Cortex | Signaling pathways regulating pluripotency of stem cells | 144 | 7 | 0.108 | 6.02 | 0.009 | Immune & Metabolic Stress |

| Brain Region | Pathway | Total | Hits | Expected | Hits/Exp. | FDR | Cluster |
|---|---|---|---|---|---|---|---|
| Frontal Cortex | TGF-beta signaling pathway | 110 | 7 | 0.082 | 6.41 | 0.002 | Immune & Metabolic Stress |
| Frontal Cortex | Thyroid hormone signaling pathway | 122 | 8 | 0.104 | 6.26 | < 0.001 | Immune & Metabolic Stress |
| Frontal Cortex | Transcriptional misregulation in cancer | 201 | 8 | 0.172 | 5.54 | 0.015 | Immune & Metabolic Stress |
| Frontal Cortex | Wnt signaling pathway | 178 | 11 | 0.209 | 5.72 | < 0.001 | Immune & Metabolic Stress |
| Frontal Cortex | Adipocytokine signaling pathway | 70 | 5 | 0.037 | 7.07 | 0.007 | Endocrine Signaling |
| Frontal Cortex | AGE-RAGE signaling pathway in diabetic complications | 101 | 6 | 0.065 | 6.54 | 0.007 | Endocrine Signaling |
| Frontal Cortex | Alcoholic liver disease | 144 | 10 | 0.154 | 6.02 | < 0.001 | Endocrine Signaling |
| Frontal Cortex | AMPK signaling pathway | 122 | 7 | 0.091 | 6.26 | 0.004 | Endocrine Signaling |
| Frontal Cortex | Autophagy - animal | 169 | 11 | 0.198 | 5.79 | < 0.001 | Endocrine Signaling |
| Frontal Cortex | Chemical carcinogenesis - reactive oxygen species | 227 | 8 | 0.194 | 5.37 | 0.028 | Endocrine Signaling |
| Frontal Cortex | C-type lectin receptor signaling pathway | 105 | 7 | 0.078 | 6.48 | 0.002 | Endocrine Signaling |
| Frontal Cortex | Endocytosis | 250 | 8 | 0.213 | 5.23 | 0.045 | Endocrine Signaling |
| Frontal Cortex | HIF-1 signaling pathway | 110 | 10 | 0.117 | 6.41 | < 0.001 | Endocrine Signaling |
| Frontal Cortex | IgSF CAM signaling | 299 | 11 | 0.351 | 4.97 | 0.007 | Endocrine Signaling |
| Frontal Cortex | IL-17 signaling pathway | 94 | 6 | 0.060 | 6.64 | 0.005 | Endocrine Signaling |
| Frontal Cortex | Insulin resistance | 109 | 5 | 0.058 | 6.43 | 0.034 | Endocrine Signaling |
| Frontal Cortex | mTOR signaling pathway | 158 | 8 | 0.135 | 5.89 | 0.004 | Endocrine Signaling |
| Frontal Cortex | Neutrophil extracellular trap formation | 195 | 7 | 0.146 | 5.59 | 0.036 | Endocrine Signaling |
| Frontal Cortex | NF-kappa B signaling pathway | 105 | 8 | 0.090 | 6.48 | < 0.001 | Endocrine Signaling |
| Frontal Cortex | Osteoclast differentiation | 142 | 10 | 0.151 | 6.05 | < 0.001 | Endocrine Signaling |
| Frontal Cortex | Pancreatic cancer | 77 | 8 | 0.066 | 6.93 | < 0.001 | Endocrine Signaling |
| Frontal Cortex | PD-L1 expression and PD-1 checkpoint pathway in cancer | 90 | 8 | 0.077 | 6.70 | < 0.001 | Endocrine Signaling |
| Frontal Cortex | RIG-I-like receptor signaling pathway | 72 | 6 | 0.046 | 7.02 | 0.001 | Endocrine Signaling |
| Frontal Cortex | T cell receptor signaling pathway | 122 | 8 | 0.104 | 6.26 | < 0.001 | Endocrine Signaling |
| Frontal Cortex | Th17 cell differentiation | 109 | 7 | 0.081 | 6.43 | 0.002 | Endocrine Signaling |
| Frontal Cortex | Type II diabetes mellitus | 47 | 4 | 0.020 | 7.64 | 0.009 | Endocrine Signaling |
| Frontal Cortex | Ubiquitin mediated proteolysis | 142 | 11 | 0.167 | 6.05 | < 0.001 | Endocrine Signaling |
| Frontal Cortex | VEGF signaling pathway | 60 | 5 | 0.032 | 7.29 | 0.004 | Endocrine Signaling |

### ADNI Acknowledgment list

ACKNOWLEDGEMENT LIST FOR ADNI PUBLICATIONS

The Data and Publications Committee, in keeping with the publication policies adopted by the ADNI Steering Committee, here provide lists for standardized acknowledgement. The list consists of two parts:

Infrastructure Investigators and Site Investigators. Infrastructure Investigators represent the names responsible for leadership and infrastructure. Site Investigators represent the names of individuals at each recruiting site. All papers, including methodological papers, should have an acknowledgement list that consists of Infrastructure Investigators plus the FULL list.

I. ADNI 1, GO, 2, 3, 4

**Part A: Leadership and Infrastructure Principal Investigator**

Michael Weiner, MD University of California, San Francisco Northern California Institute for Research and Education
ATRI PI and Director of Coordinating Center Clinical Core
Paul Aisen, MD University of Southern California
Ronald Petersen, MD, PhD Mayo Clinic, Rochester (co-PI of of Clinical Core)

Executive Committee
Michael Weiner, MD University of California, San Francisco
Paul Aisen, MD University of Southern California
Ronald Petersen, MD, PhD Mayo Clinic, Rochester
Clifford R. Jack, Jr., MD Mayo Clinic, Rochester
William Jagust, MD University of California, Berkeley
Susan Landau, PhD University of California, Berkeley
Monica Rivera-Mindt, PhD Fordham University; Mt. Sinai Medical Center Ozioma Okonkwo, PhD University of Wisconsin
Leslie M. Shaw, PhD University of Pennsylvania
Edward B. Lee, MD, PhD University of Pennsylvania
Arthur W. Toga, PhD University of California, Los Angeles
Laurel Beckett, PhD University of California, Davis
Danielle Harvey, PhD University of California, Davis
Robert C. Green, MD, MPH Boston University
Andrew J. Saykin, PsyD Indiana University
Kwangsik Nho, PhD Indiana University
Richard J. Perrin, MD, PhD Washington University St. Louis
Duygu Tosun, PhD University of California, San Francisco

ADNI 4 Private Partner Scientific Board (PPSB) Convened by Alzheimer's Association

Pallavi Sachdev, PhD Eisai (Chair, 2023-2024)

Data and Publication Committee (DPC)
Robert C. Green, MD, MPH Harvard University (Chair)
Erin Drake Harvard University

Resource Allocation Review Committee
Tom Montine, MD, PhD University of Washington (Chair)
Cat Conti, BA Northern California Institute for Research and Education

Administrative Core Leaders and Key Personnel
Michael W. Weiner, MD University of California, San Francisco
Rachel Nosheny, PhD University of California, San Francisco
Diana Truran Sacrey Northern California Institute for Research and Education
Juliet Fockler University of California, San Francisco
Melanie J. Miller, PhD Northern California Institute for Research and Education
Catherine (Cat) Conti Northern California Institute for Research and Education
Winnie Kwang, MA University of California, San Francisco
Chengshi Jin, PhD University of California, San Francisco
Adam Diaz, MSNorthern California Institute for Research and Education
Miriam Ashford, PhD Northern California Institute for Research and Education
Derek Flenniken Northern California Institute for Research and Education
Adrienne Kormos Northern California Institute for Research and Education
Clinical Core Leaders and Key Personnel
Ronald Petersen, MD, PhD Mayo Clinic, Rochester (Core PI)
Paul Aisen, MDUniversity of Southern California (Core PI)
Michael Rafii, MD, PhD University of Southern California
Rema Raman, PhD University of Southern California
Gustavo Jimenez, MBS University of Southern California
Michael Donohue, PhD University of Southern California
Jennifer Salazar, MBS University of Southern California
Andrea Fidell, MPH University of Southern California
Virginia Boatwright, BSUniversity of Southern California
Justin Robison, MS University of Southern California
Caileigh Zimmerman, MS University of Southern California
Yuliana Cabrera, BS University of Southern California

Sarah Walter, MSc University of Southern California
Taylor Clanton, MPH University of Southern California
Elizabeth Shaffer, BS University of Southern California
Caitlin Webb, BA University of Southern California
Lindsey Hergesheimer, BS University of Southern California
Stephanie Smith, BS University of Southern California
Sheila Ogwang, MPH University of Southern California
Olusegun Adegoke, MSc University of Southern California
Payam Mahboubi, MPH University of Southern California
Jeremy Pizzola, BA University of Southern California
Cecily Jenkins, PhD University of Southern California

Biostatistics Core Leaders and Key Personnel
Laurel Beckett, PhD University of California, Davis (Core PI)
Danielle Harvey, PhD University of California, Davis (Core PI)
Michael Donohue, PhD University of Southern California
Naomi Saito, MS University of California, Davis
Adam Diaz, MS Northern California Institute for Research and Education
Kedir Adem Hussen, MS University of Southern California

Engagement Core Leaders and Key Personnel
Ozioma Okonkwo, PhD University of Wisconsin (Core-PI)
Monica Rivera-Mindt, PhD Fordham University; Mt. Sinai (Core-PI)
Hannatu Amaza University of Wisconsin
Mai Seng Thao University of Wisconsin
Shaniya Parkins Mt. Sinai
Omobolanle Ayo, MBChB, MPH Mt. Sinai
Matt Glittenberg University of Wisconsin
Isabella Hoang University of Wisconsin
Kaori Kubo Germano, PhD Fordham University
Joe Strong, PhD University of Wisconsin
Trinity Weisensel University of Wisconsin
Fabiola Magana University of Wisconsin
Lisa Thomas University of Wisconsin
Vanessa Guzman, PhD Mt. Sinai
Adeyinka Ajayi, MBBS, MPH Mt. Sinai
Joseph Di Benedetto, LMSW Mt. Sinai
Sandra Talavera, MSW Fordham University

MRI Core Leaders and Key Personnel
Clifford R. Jack, Jr., MD Mayo Clinic, Rochester (Core PI)

Joel Felmlee, PhD Mayo Clinic, Rochester
Nick C. Fox, MD University College London
Paul Thompson, PhD UCLA School of Medicine
Charles DeCarli, MD University of California, Davis Arvin Forghanian-Arani, PhD Mayo Clinic, Rochester
Bret Borowski, RTR Mayo Clinic, Rochester
Calvin Reyes Mayo Clinic, Rochester
Caitie Hedberg Mayo Clinic, Rochester
Chad Ward Mayo Clinic, Rochester Christopher Schwarz, PhD Mayo Clinic, Rochester Denise Reyes Mayo Clinic, Rochester
Jeff Gunter, PhD Mayo Clinic, Rochester
John Moore-Weiss, PhDMayo Clinic, Rochester
Kejal Kantarci, MD Mayo Clinic, Rochester
Leonard Matoush Mayo Clinic, Rochester
Matthew Senjem, MS Mayo Clinic, Rochester
Prashanthi Vemuri, PhDMayo Clinic, Rochester
Robert Reid, PhD Mayo Clinic, Rochester
Ian Malone, PhD University College London
Sophia I. Thomopoulos, BS University of Southern California School of Medicine Talia M. Nir, PhD University of Southern California School of Medicine Neda Jahanshad, PhD University of Southern California School of Medicine Alexander Knaack, MS University of California, Davis
Evan Fletcher, PhD University of California, Davis
Danielle Harvey, PhD University of California, Davis
Duygu Tosun-Turgut, PhD University of California, San Francisco
Stephanie Rossi Chen, BA. Northern California Institute for Research and Education Mark Choe, BS Northern California Institute for Research and Education
Karen CrawfordUniversity of Southern California School of Medicine Paul A. Yushkevich, PhD University of Pennsylvania
Sandhitsu Das, PhD University of Pennsylvania

PET Core Leaders and Key Personnel
William Jagust, MD University of California, Berkeley (Core PI)
Susan Landau, PhD University of California, Berkeley (Core PI)
Robert A. Koeppe, PhD University of Michigan
Gil Rabinovici, MD University of California San Francisco
Victor Villemagne, MD University of Pittsburgh
Brian LoPresti, MSNE University of Pittsburgh

Neuropathology Core Leaders and Key Personnel
Richard J. Perrin, MD, PhD Washington University St. Louis (Core PI) John Morris, MD Washington University St. Louis
Erin Franklin, MS Washington University St. Louis

Haley Bernhardt, BA, R. EEG T. Washington University St. Louis Nigel J. Cairns, PhD, MRCPath Washington University St. Louis Lisa Taylor-Reinwald, BA, HTL (ASCP) Washington University St. Louis

Biomarkers Core Leader and Key Personnel
Leslie Shaw, PhD UPenn School of Medicine (Core PI) Edward B. Lee, MD, PhD University of Pennsylvania (Core PI) Virginia M.Y. Lee, PhD, MBA UPenn School of Medicine Magdalena Korecka, PhD UPenn School of Medicine Magdalena Brylska, MS UPenn School of Medicine
Yang Wan, MS UPenn School of Medicine
J.Q. Trojanowki, MD, PhD* UPenn School of Medicine (*former Core PI, deceased)

Informatics Core Leader and Key Personnel
Arthur W. Toga, PhD University of Southern California (Core PI) Karen Crawford, MLIS University of Southern California
Scott Neu, PhD University of Southern California

Genetics Core Leader and Key Personnel
Andrew J. Saykin, PsyDIndiana University School of Medicine (Core PI) Kwangsik Nho, PhD Indiana University School of Medicine (Core PI) Tatiana M. Foroud, PhD Indiana University School of Medicine (Dir. NCRAD) Taeho Jo, PhD Indiana University School of Medicine
Shannon L. Risacher, PhD Indiana University School of Medicine Hannah Craft, MPH Indiana University School of Medicine Liana G. Apostolova, MDIndiana University School of Medicine
Kelly Nudelman, PhD NCRAD/Indiana University School of Medicine Kelley Faber, MS, CCRC NCRAD/Indiana University School of Medicine Zoë Potter, BA, CCRP NCRAD/Indiana University School of Medicine Kaci Lacy, MPH, CCRP NCRAD/Indiana University School of Medicine
Rima Kaddurah-Daouk, PhD Duke University/AD Metabolomics Consortium Li Shen, PhD University of Pennsylvania
ADNI4 Amyloid PET Visual Read Team
David Soleimani-Meigooni, MD University of California, San Francisco Renaud La Joie, PhD University of California, San Francisco Konstantinos Chiotis, MD, PhD University of California, San Francisco Maison Abu Raya, MD University of California, San Francisco Agathe Vrillon, MD, PhD University of California, San Francisco Charles Windon, MD University of California, San Francisco Julien Lagarde, MD, PhD University of California, San Francisco Zoe Lin University of California, San Francisco
Aidyn Rose Hills University of California, San Francisco

ADNI4 Amyloid Disclosure Team
Jason Karlawish, MD University of Pennsylvania
Claire Erickson, PhD University of Pennsylvania
Joshua Grill PhD University of California, Irvine
Emily Largent PhD University of Pennsylvania
Kristin Harkins MPH University of Pennsylvania

Early Project Development
Michael W. Weiner, MD UCSF/NCIRE Leon Thal, MD – Past Investigator
Zaven Kachaturian, PhD Khachaturian, Radebaugh & Associates (KRA), Inc Richard Frank, MD, PhD General Electric
Peter J. Snyder, PhD University of Connecticut Alzheimer's Association's Ronald and Nancy Reagan's Research Institute

NIA
Neil Buckholtz, PhD National Institute on Aging
John K. Hsiao, MD National Institute on Aging
Laurie Ryan, PhD National Institute on Aging
Susan Molchan, PhD National Institute on Aging/National Institutes of Health

ADNI External Scientific Advisory Board (SAB)
Zaven Khachaturian, PhD Prevent Alzheimer's Disease 2020 (Chair) Maria Carrillo, PhD Alzheimer's Association
William Potter, MD National Institute of Mental Health
Lisa Barnes, PhD Rush University
Marie Bernard, MD NIA
Hector González University of California, San Diego
Carole Ho Denali Therapeutics
John K. Hsiao, MD NIH
Jonathan Jackson, PhD Massachusetts General Hospital
Eliezer Masliah, MD NIA
Donna Masterman, MD Biogen
Ozioma Okonkwo, PhD University of Wisconsin, Madison Richard Perrin, MD, PhD Washington University St. Louis Laurie Ryan, PhD NIA
Nina Silverberg, PhD NIA

Part B: Investigators By Site

### Oregon Health and Science University:

Lisa Silbert, MD Jeffrey Kaye, MD
Sylvia White (Salazar), ND Aimee Pierce, MD
Amy Thomas, BSN, RN Tera Clay
Daniel Schwartz, BA Gillian Devereux, RN, MPH Janet "Janae" Taylor Jennifer Ryan, ND, MS Mike Nguyen
Madison DeCapo, BS Yanan Shang, MD
University of Southern California:
Lon Schneider, MD Cynthia Munoz, MA Diana Ferman, PA Carlota Conant, BS Katherin Martin Kristin Oleary
Sonia Pawluczyk, MD Elizabeth Trejo
Karen Dagerman Liberty Teodoro, RN Mauricio Becerra Madiha Fairooz, BS Sonia Garrison, MSsc Julia Boudreau, MS Yair Avila, BA
University of California--San Diego:
James Brewer, MD, PhD Aaron Jacobson
Antonio Gama Chi Kim
Emily Little, MPH Jennifer Frascino Nichol Ferng
Socorro Trujillo, MPH

**University of Michigan:** Judith Heidebrink, MD Robert Koeppe, PhD Steven MacDonald, MD

Dariya Malyarenko, Ph.D.
Jaimie Ziolkowski, MA, BS, TLLP James O'Connor, MS, RT (R)(MR) Nicole Robert
Suzan Lowe Virginia Rogers

**Mayo Clinic, Rochester:** Ronald Petersen, MD, Ph.D. Barbara Hackenmiller Bradley Boeve, MD Colleen Albers, RN
Connie Kreuger David Jones, MD David Knopman, MD
Hugo Botha, MB, Ch.B. Jessica Magnuson
Jonathan Graff-Radford, MD Kerry Crawley, BSW, CCRP Michael Schumacher, CNMT Sanna McKinzie, MS
Steven Smith, MS Tascha Helland, BS Val Lowe, MD
Vijay Ramanan, MD, PhD

Baylor College of Medicine:
Valory Pavlik, PhD Jacob Faircloth, BS Jeffrey Bishop, PA Jessica Nath
Maria Chaudhary, MAP Maria Kataki, PhD, MD Melissa Yu, MD, FAAN Nathiel Pacini, MA Randall Barker
Regan Brooks, BA Ruchi Aggarwal, MD

Columbia University Medical Center:
Lawrence Honig, MD, Ph.D. Yaakov Stern, PhD

Akiva Mintz, MD Jonathan Cordona, ARRT Michelle Hernandez

Washington University, St. Louis:
Justin Long, MD Abbey Arnold, NP Alex Groves
Anna Middleton, RN Blake Vogler
Cierra McCurry Connie Mayo, RN Cyrus Raji, MD, PhD
Fatima S. Amtashar, BS Heather Klemp, MSW
Heather Nicole Elmore, RN, MSN, ANP-BC, CCRP James Ruszkiewicz, CNMT
Jasmina Kusuran Jasmine Stewart
Jennifer Horenkamp, RN, BSN Julia Greeson, MS
Kara Wever, MA Katie Vo, MD Kelly Larkin, RN Lesley Rao, MD
Lisa Schoolcraft, BFA Lora Gallagher
Madeline Paczynski, BS, PA-C Maureen McMillan
Michael Holt, MSW Nicole Gagliano, BS, RT Rachel Henson, MS Renee LaBarge
Robert Swarm, MD Sarah Munie, BSN, RN Serena Cepeda, BS
Stacey Winterton, BSN, RN Stephen Hegedus
TaNisha Wilson Tanya Harte, FNP-BC Zach Bonacorsi

University of Alabama Birmingham:
David Geldmacher, MD

Amber Watkins, RN Brandi Barger, BSRT Bryan Smelser, MD Charna Bates, MA
Cynthia Stover, PENDING Emily McKinley,
Gregory Ikner, MA Haley Hendrix,
Harold Matthew Cooper, MSN, CRNP, NPC Jennifer Mahaffey,
Lindsey Booth Robbins, MSN, CRNP, PNP-C Loren Brown Ashley, RN, BSN
Marissa Natelson-Love, MD Princess Carter, RN Veronika Solomon,
Mount Sinai School of Medicine:
Hillel Grossman, MD Alexandra Groome, BA Allison Ardolino, MA
Anthony Kaplan, ARRT, CNMT Faye Sheppard, BS
Genesis Burgos-Rivera, BA Gina Garcia-Camilo, MD Joanne Lim, MA
Judith Neugroschl, MD Kimberly Jackson, BS Kirsten Evans, BS Laili Soleimani, MD Mary Sano, Ph.D.
Nasrin Ghesani, MD Sarah Binder, BS
Xiomara Mendoza Apuango, BS

Rush University Medical Center:
Ajay Sood, MD, PhD Amelia Troutman, MA
Kimberly Blanchard, APRN, DNP, NP-C Arlene Richards,
Grace Nelson, BA
Kirsten Hendrickson, RN, MSN Erin Yurko,
Jamie Plenge, BS Victoria Rufo, MS Raj Shah, MD

**Wein Center**: Ranjan Duara, MD Brendan Lynch, CRT Cesar Chirinos, PsyD
Christine Dittrich, CRT Debbie Campbell Diego Mejia, CRT Gilberto Perez, CRT Helena Colvee, BS
Joanna Gonzalez, PsyD Josalen Gondrez, MS Joshua Knaack
Mara Acevedo
Maria Cereijo, APRN Maria Greig-Custo, MD Michelle Villar, BS Morris Wishnia
Sheryl Detling Warren Barker, MS

**Johns Hopkins University:** Marilyn Albert, Ph.D. Abhay Moghekar
Barbara Rodzon Corey Demsky Gregory Pontone, MD Jim Pekar
Leonie Farrington, CNRN Martin Pomper
Nicole Johnson Tolulope Alo

**New York University:** Martin Sadowski, MD, PhD Anaztasia Ulysse, BA Arjun Masurkar
Brittany Marti David Mossa, R.T Emilie Geesey Emily Petrocca, NP Evan Schulze, PhD Jennifer Wong
Joseph Boonsiri
Sunnie Kenowsky, DVM

Tatianne Martinez, NP Veronica Briglall

Duke University Medical Center:
P. Murali Doraiswamy, MD, MBBS Adaora Nwosu
Alisa Adhikari, BS Cammie Hellegers, MA Jeffrey Petrella
Olga James, MD Terence Wong Thomas Hawk
**University of Pennsylvania:** Sanjeev Vaishnavi, MD, PhD Hannah McCoubrey, BA
Ilya Nasrallah, MD, PhD Rachel Rovere, BA Jeffrey Maneval, MD Elizabeth Robinson, MA Francisco Rivera, MS Jade Uffelman, BS Martha Combs, BS, MS Patricia O'Donnell
Sara Manning, MD

**University of Kentucky:** Richard King, MD Alayne Nieto, BSN, RN Amanda Glueck, PhD Anjana Mandal
Audrie Swain
Bethanie Gamble, PhD, RN Beverly Meacham, RT(R) (MR) Denece Forenback, RN
Dorothy Ross, CCRP Elizabeth Cheatham Ellen Hartman
Gary Cornell Jordan Harp, PhD Laura Ashe Laura Goins Linda Watts, RN Morgan Yazell Prabin Mandal

Regan Buckler, BSN, RN Sylvia Vincent
Triana Rudd

University of Pittsburgh:
Oscar Lopez, MD Ann Arlene Malia Caitlin Chiado, CRNP Cary Zik
James Ruszkiewicz, CNMT Kathleen Savage
Linda Fenice MaryAnn Oakley, MA Paige C Tacey, M.Ed.
Sarah Berman, MD, PhD Sarah Bowser, CRNP Stephen Hegedus Xanthia Saganis

University of Rochester Medical Center:
Anton Porsteinsson, MD Abigail Mathewson, RN, BSN Asa Widman, BA
Bridget Holvey, BS Emily Clark, DO Esmeralda Morales, MS Iris Young, PA-C
James Ruszkiewicz, CNMT
Kevin Hopkins, BS, CNMT, LNMT Kimberly Martin, RN, BSN
Nancy Kowalski, RN, MS Rebecca Hunt, BS Roberta Calzavara, PhD
Russell Kurvach, BS, CCRP Stephen D'Ambrosio, PA-C, MPAS
University of California, Irvine:
Gaby Thai, MD
Beatriz Vides, RN, MSN Brigit Lieb, ARRT/CRT
Catherine McAdams-Ortiz, MSN, RN, A/GNP Cyndy Toso
Ivan Mares, BS Kathryn Moorlach

Luter Liu
Maria Corona, PhD Mary Nguyen, BA
Melanie Tallakson, DNP, FNP-C Michelle McDonnell, PhD Milagros Rangel, BS
Neetha Basheer, MD, MBBS Patricia Place, BA
Romina Romero, PhD Steven Tam, MD

University of Texas Southwestern Medical School:
Trung Nguyen, MD, PhD Abey Thomas, ARRT Alexander (Alex) Frolov, MD Alka Khera, MD
Amy Browning, BA (Pending) Brendan Kelley (031), MD Courtney Dawson, RT(R) Dana Mathews, MD, Ph.D. Elaine Most, MS (Pending) Elizeva (Ellie) Phillips, CNMT Lynn Nguyen
Maribel Nunez Matalin Miller, MS Matthew R. Jones, MA
Natalie Martinez, MSN, RN, FNP-BC Rebecca Logan, PA-C
Roderick McColl Sari Pham
Tiffani Fox, MBA, MS Tracey Moore, BA

**Emory University:** Allan Levey, MD, PhD Abby Brown, NP Andrea Kippels, NP
Ashton Ellison, BSPH, ABA Casie Lyons
Chadwick Hales, MD, PhD Cindy Parry, BFA Courtney Williams Elizabeth McCorkle, BS Guy Harris, BA

Heather Rose, BSN Inara Jooma, BS
Jahmila Al-Amin, MS, BS James Lah, MD, PhD James Webster, BS
Jessica Swiniarski, MPH, BS Latasha Chapman, BS
Laura Donnelly, MPH Lauren Mariotti
Mary Locke, BS Phyllis Vaughn, BSN
Rachael Penn, BSN, RN Sallie Carpentier, RN, BSN
Samira Yeboah, BMSc, R.T.(R) (MR) Sarah Basadre, BMSc, ARRT(R)(MR) Sarah Malakauskas, MS
Stefka Lyron, NP Tara Villinger, NP Terra Burney
University of Kansas, Medical Center:
Jeffrey Burns, MD, MS Ala Abusalim, PA-C Alexandra Dahlgren, BS Alexandria Montero, RN
Anne Arthur, BSN, MS, ANP-BC Heather Dooly, BS
Katelynn Kreszyn, APRN Katherine Berner, BS Lindsey Gillen, APRN Maria Scanlan, BA Mercedes Madison, BS Nicole Mathis
Phyllis Switzer Ryan Townley, MD
Samantha Fikru, APRN, MSN, FNP-C Samantha Sullivan, MSW
Ella Wright, BS
**University of California, Los Angeles**: Maryam Beigi, MD
Anthony Daley Ashley Ko Brittney Luong Glen Nyborg

Jessica Morales Kelly Durbin, PhD Lauren Garcia Leila Parand Lorena Macias
Lorena Monserratt, PhD Maya Farchi
Pauline Wu, DO Robert Hernandez Thao Rodriguez, NP
Mayo Clinic, Jacksonville:
Neill Graff-Radford, MD, MBBCH, FRCP A'llana Marolt, BS
Anton Thomas, BS Deborah Aloszka Ercilia Moncayo, BS Erin Westerhold, RT Gregory Day, MD
Kandise Chrestensen, BS Mary Imhansiemhonehi, BS Sanna McKinzie, MS Sochenda Stephens, CCRP
Sylvia Grant, CCRC
**Indiana University:** Jared Brosch, MD Amy Perkins, CCRP Aubree Saunders, BS
Debra Silberberg Kovac, BS Heather Polson, CNMT Isabell Mwaura, BS Kassandra Mejia, BS Katherine Britt, BS
Kathy King, RN Kayla Nichols, BS Kayley Lawrence, BA Lisa Rankin, BSW Martin Farlow, MD
Patricia Wiesenauer, MS Robert Bryant, BS
Scott Herring, RN Sheryl Lynch, RN Skylar Wilson Traci Day

William Korst

Yale University School of Medicine:
Christopher van Dyck, MD Adam Mecca, MD, PhD Alyssa Miller, BS
Amanda Brennan, LMSE, MSW Amber Khan, MD
Audrey Ruan
Carol Gunnoud, AS Chelsea Mendonca, MD
Danielle Raynes-Goldfinger, BS Elaheh Salardini, MD
Elisa Hidalgo, MS, CNMT, EMT, RT (CT) Emma Cooper, BA
Erawadi Singh, DO Erin Murphy, BS
Jeanine May, APRN, MSN, MHP, CCRP
Jesse Stanhope, BS Jessica Lam, BSE Julia Waszak, BS Kimberly Nelsen, BA Kimberly Sacaza, BS
Mayer Joshua Hasbani, MD Meghan Donahue, BA Ming-Kai Chen, MD, PhD Nicole Barcelos, MS, MA
Paul Eigenberger, MD Robin Bonomi, MD
Ryan O'Dell, MD, PhD Sarah Jefferson, MD Siddharth Khasnavis, MD Stephen Smilowitz, MD
Susan DeStefano, APRN, MSN Susan Good, APRN
Terry Camarro, RT, RN, MRI, APRT Vanessa Clayton, BS
Yanis Cavrel, BA YuQuan "Oliver" Lu
McGill University, Montreal-Jewish General Hospital:
Howard Chertkow, MD Howard Bergman, MD Chris Hosein, M.Ed

Sunnybrook Health Sciences, Ontario:
Sandra Black, MD Anish Kapadia, MD Aparna Bhan
Benjamin Lam, MD, FRCP(c) Christopher Scott, BSc Gillian Gabriel, MA
Jennifer Bray, BA, BSW, MSW Ljubica Zotovic, MD
Maria Samira Gutierrez Mario Masellis
Marjan Farshadi, MD Maurylette Gui, Psych BSc Meghan Mitchell, BSc Rebecca Taylor
Ruby Endre, M.R.T Zhala Taghi-Zada
University of British Columbia Clinic for AD & Related Disorders
Robin Hsiung, MD Carolyn English Ellen Kim, BA Eugene Yau
Haley Tong
Laura Barlow, RTR/RTMR Lauren Jennings
Michele Assaly Paula Nunes, PhD Tahlee Marian

Cognitive Neurology St. Joseph's Ontario:
Andrew Kertesz, MD John Rogers, MD Dick Trost, PhD
Cleveland Clinic Lou Ruvo Center for Brain Health
Dylan Wint, MD Charles Bernick, MD Donna Munic, PhD
Northwestern University:
Ian Grant, MD Aaliyah Korkoyah, BS Ali Raja
Allison Lapins, MD

Caila Ryan, MS Jelena Pejic
Kailey Basham, BS Leena Lukose, BS Loreece Haddad, MS
Lucas Quinlan, BS, MLS (ASCP) Nathaniel Houghtaling
Premiere Research Inst (Palm Beach Neurology):
Carl Sadowsky MD Walter Martinez MD Teresa Villena MD
Georgetown University Medical Center:
Brigid Reynolds, NP Angelica Forero, MS Carolyn Ward, MSPH Emma Brennan, BS Esteban Figueroa
Giuseppe Esposito, MD Jessica Mallory
Kathleen Johnson, RN, NP Kathryn Turner, BSN Katie Seidenberg
Kelly McCann, BA Margaret Bassett, NP Melanie Chadwick, NP
Raymond Scott Turner, MD, PhD Robin Bean, RT
Saurabh Sharma, MD
Brigham and Women's Hospital:
Gad Marshall, MD Aferdita Haviari, BA Alison Pietras, PA-C, ACP Bradley Wallace, BS Catherine Munro, PhD
Gladiliz Rivera-Delpin, MA Hadley Hustead, BS Isabella Levesque
Jennifer Ramirez, BA Karen Nolan, BS, RT (MR)
Kirsten Glennon, RN, CNRN Mariana Palou, BA
Michael Erkkinen, MD

Nicole DaSilva
Pamela Friedman, Psy. D Regina M. Silver, RN Ricardo Salazar, MD Roxxanne Polleys, AA Scott McGinnis (094), MD Seth Gale, MD
Tia Hall, BS Tuan Luu

Stanford University:
Steven Chao, MD Emmeline Lin, BS Jaila Coleman, BA
Kevin Epperson, RT(R)(MR) Minal Vasanawala
Banner Sun Health Research Institute
Alireza Atri, MD, PhD Amy Rangel
Brittani Evans Candy Monarrez Carol Cline, LMSW
Carolyn Liebsack, RN, BSN, CCRC Daniel Bandy
Danielle Goldfarb, MD Debbie Intorcia Jennifer Olgin
Kelly Clark
Kelsey King, CCRP Kylee York
Marina Reade, RN, FNP-C Michael Callan
Michael Glass
Michaela Johnson, G-ACNP, BC Michele Gutierrez
Molly Goddard
Nadira Trncic, MD, PhD Parichita Choudhury, MD Priscilla Reyes
Serena Lowery Shaundra Hall Sonia Olgin
Stephanie de Santiago, RN, NP

Boston University:
Michael Alosco, PhD Alyssa Ton, BS
Amanda Jimenez, MS, EMT-B, CPT Andrew Ellison, MR Technologist Anh Tran, RN
Brandon Anderson, RT(N), CNMT Della Carter, MS
Donna Veronelli, RTN, CNMT Steven Lenio, MD
Eric Steinberg, RN, MSN, CNP Jesse Mez, MD, MS
Jason Weller, MD Jennifer Johns, RN Jesse Mez, MD, MS Jessica Harkins, CNMT Alexa Puleio, MS
Ina Hoti, BS
Jane Mwicigi, MBChB., MPH Alexa Puleio, MS
Michael Alosco, PhD Olivia Schultz, BA Mona Lauture, RN Eric Steinberg Ridiane Denis, RN Ronald Killiany, PhD Sarab Singh, CNMT Steven Lenio, MD Wendy Qiu, MD, PhD Ycar Devis, MPH
Howard University:
Thomas Obisesan, MD, MPH Andrew Stone, MS
Debra Ordor, RN, BSN Ifreke Udodong, CRNP
Immaculata Okonkwo, DNP, MSN, APRN, FNP-BC Javed Khan, MD
Jillian Turner, BS, MS Kyliah Hughes, BS, RMA Oshoze Kadiri, MPH
Case Western Reserve University:
Charles Duffy, MD, PhD

Ariana Moss
Katherine Stapleton, LPN Maria Toth (fmr Gross), RN Marianne Sanders, BSN, RN Martin Ayres
Melissa Hamski Parianne Fatica, CCRC Paula Ogrocki, PhD Sarah Ash
Stacy Pot
University of California, Davis Sacramento :
Doris Chen, MD Andres Soto
Costin Tanase, PhD David Bissig, MD, PhD Hafsanoor Vanya, BA
Heather Russell (116), CNMT Hitesh Patel, CNMT Hongzheng Zhang, CCRP Kelly Wallace, CCRP
Kristi Ayers, BS Maria Gallegos, BS Martha Forloines, PhD Meghan Sinn
Queennie Majorie S Kahulugan, CCRC Richard Isip, RT (R)(N)(CT)
Sandra Calderon, MS, RN, FMP-C Talia Hamm, BA, CCRP
**Parkwood Hospital:** Michael Borrie, MD T-Y Lee, PhD
Dr Rob Bartha, PhD
**University of Wisconsin:** Sterling Johnson, PhD Sanjay Asthana, MD Cynthia M. Carlsson, MD

Banner Alzheimer's Institute:
Allison Perrin, MD Pierre Tariot, MD Adam Fleisher, MD Stephanie Reeder, BA

**Dent Neurologic Institute** Horacio Capote, MD Allison Emborsky
Anna Mattle, PharmD, MS Bela Ajtai, MD
Benjamin Wagner, PA-C Bennett Myers
Daryn Slazyk
Delaney Fragale, PA-C Erin Fransen, PA Heather Macnamara Jonathan Falletta, PA-C Joseph Hirtreiter, RN Laszlo Mechtler, MD Megan King
Michael Asbach, RPA-C
Michelle Rainka, Pharm. D., CCRP Richard Zawislak, NP
Scott Wisniewski Stephanie O'Malley, PA-C Tatiana Jimenez-Knight Todd Peehler
Traci Aladeen, PharmD Vernice Bates
Violet Wenner Wisam Elmalik, MD
Ohio State University:
Douglas W. Scharre, MD Arun Ramamurthy, MD Soumya Bouchachi, MD
Maria Kataki, MD, PhD - Past Investigator Rawan Tarawneh, MD - Past Investigator Brendan Kelley, MD - Past Investigator

**Albany Medical College:** Dzintra Celmins, MD Alicia Leader
Chris Figueroa Heather Bauerle, NP Katlynn Patterson Michael Reposa Steven Presto
Tuba Ahmed

Wendy Stewart

Hartford Hosp, Olin Neuropsychiatry Research Center:
Godfrey D. Pearlson MD Karen Blank, MD
Karen Anderson, RN
Dartmouth-Hitchcock Medical Center:
Robert B. Santulli, MD Eben S. Schwartz, PhD

Wake Forest University Health Sciences:
Jeff Williamson, MD, MHS, FACP Alicia Jessup, RN
Andrea Williams Crystal Duncan
Abigail O'Connell, APRN, FNP-C Karen Gagnon
Ezequiel Zamora James Bateman
Freda Crawford, CNMT Deb Thompson
Eboni Walker Jennifer Rowell Mikell White, MHA
Phillip "Hunter" Ledford Sarah Bohlman, MSL Susan Henkle, RN Joseph Bottoms, CNMT
Lena Moretz, RT(R) CT (MR) Bevan Hoover, BS
Michael Shannon Samantha Rogers, PA-C Wendy Baker
William Harrison, MD

**Rhode Island Hospital:** Chuang-Kuo Wu, MD Alexis DeMarco, BS
Ava Stipanovich, BS, ScM
Daniel Arcuri, CNMT, RT(N)(CT) Jan Clark, RN, BSN, CCRC, CSNT
Jennifer Davis, PhD Kerstin Doyon, RN, BSN

Marie Amoyaw, BA
Mauro Veras Acosta, PENDING, BS Ronald Bailey, RT-R, CNMT
Scott Warren, MD Terry Fogerty
Victoria Sanborn, PhD

Butler Hospital
Meghan Riddle, MD Stephen Salloway, MD, MS Paul Malloy, PhD
Stephen Correia, PhD

University of California San Francisco
Charles Windon, MD Morgan Blackburn Howard J. Rosen, MD Bruce L. Miller, MD

University of South Florida, Byrd Institute
Amanda Smith, MD Ijeoma Mba, MBA, MPH Jenny Echevarria
Juris Janavs

**University of Chicago** Emily Roglaski, PhD Meagan Yong Rebecca Devine
Eastern Virginia Medical School
Hamid Okhravi, MD

Charter Health Research Services
Edgardo Rivera, MD Teresa Kalowsky Caroline Smith Christina Rosario

Houston Methodist Neurological Institute
Joseph Masdeu, MD, PhD Richard Le, PharmD Maushami Gurung

**Barrow Neurological Institute** Marwan Sabbagh, MD Angelica Garcia
Micah Ellis Slaughter Nadeen Elayan Skieff Acothley
**Nathan Kline Institute** Nunzio Pomara, MD Raymundo Hernando Vita Pomara
Chelsea Reichert

Ralph Johnson Veterans Administration Health Care Services
Olga Brawman-Mintzer, MD Allison Acree
Arthur Williams Campbell Long Rebecca Long

Vanderbilt University Medical Center
Paul Newhouse, MD Sydni Jenee Hill Amy Boegel
University of Texas Health, San Antonio
Sudha Seshadri, MD Amy Saklad
Floyd Jones

Rutgers University
William Hu, MD, PhD
V. Sotelo
Gonzalez & Aswad Health Services
Yaneicy Gonazalez Rojas, MD
Medical University South Carolina
Jacobo Mintzer, MD, MBA Crystal Flynn Longmire, PhD Kenneth Spicer, MD, PhD

### Supplemental Figure Legends

**Fig. S1. Correlations of miR-181a, b, c, d.** Levels of miR-181a, b, c, d were measured and correlations of each measurement from the same sample were performed.

**Fig. S2. Relationships between APP and neprilysin mRNA levels.**

**Fig. S3. MAPT 3' and 5'-UTR activity assay.** A-B) MAPT 3' and 5'- UTR reporter clone constructs. C) MAPT 3'-UTR activities with miR-181 transfection. D-E) MAPT 5'-UTR activities in both HeLa and differentiated SK-N-SH cells.

**Fig. S4. PAM clustering of KEGG pathways by shared members for hippocampus.** Shared members of KEGG pathways enriched in the frontal cortex network were used to estimate Jaccard distances and clustered (k = 5) by partitioning around medoids. Cluster names are subjective, based on member pathway names. A) Neurodegeneration & Proteostasis (Fig 8F). B) Immune & Stress

Signaling. C) Cancer & Cell Fate. D) Endocrine & Signal Transduction. E) Synaptic & Neuroactive Signaling. Order is in cluster medoid Jaccard distance from cluster A.

**Fig. S5. PAM clustering of KEGG pathways by shared members for frontal cortex.** Shared members of KEGG pathways enriched in the frontal cortex network were used to estimate Jaccard distances and clustered (k = 5) by partitioning around medoids. Cluster names are subjective, based on member pathway names. A) Neurodegeneration & Apoptosis (Fig 8H). B) Synaptic Signaling. C) Cancer & Cell Fate. D) Immune & Stress Signaling. E) Endocrine Signaling. Order is in cluster medoid Jaccard distance from cluster A.

**Fig. S6. Cell type distribution of miR-181d.** A) Measurement of mir-181d mRNA by cell type.

# S1

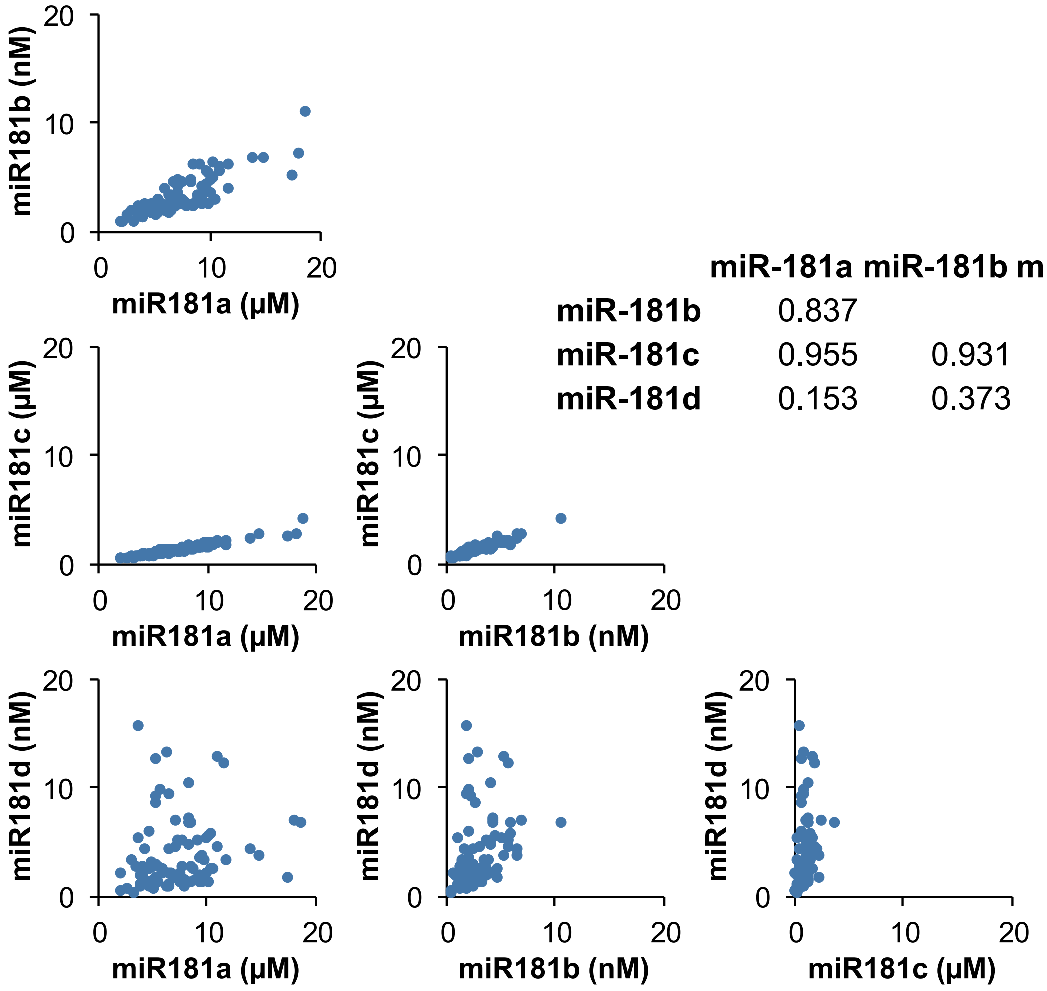


| | miR-181a | miR-181b | miR-181c |
|---|---|---|---|
| **miR-181b** | 0.837 | | |
| **miR-181c** | 0.955 | 0.931 | |
| **miR-181d** | 0.153 | 0.373 | 0.274 |

# S2

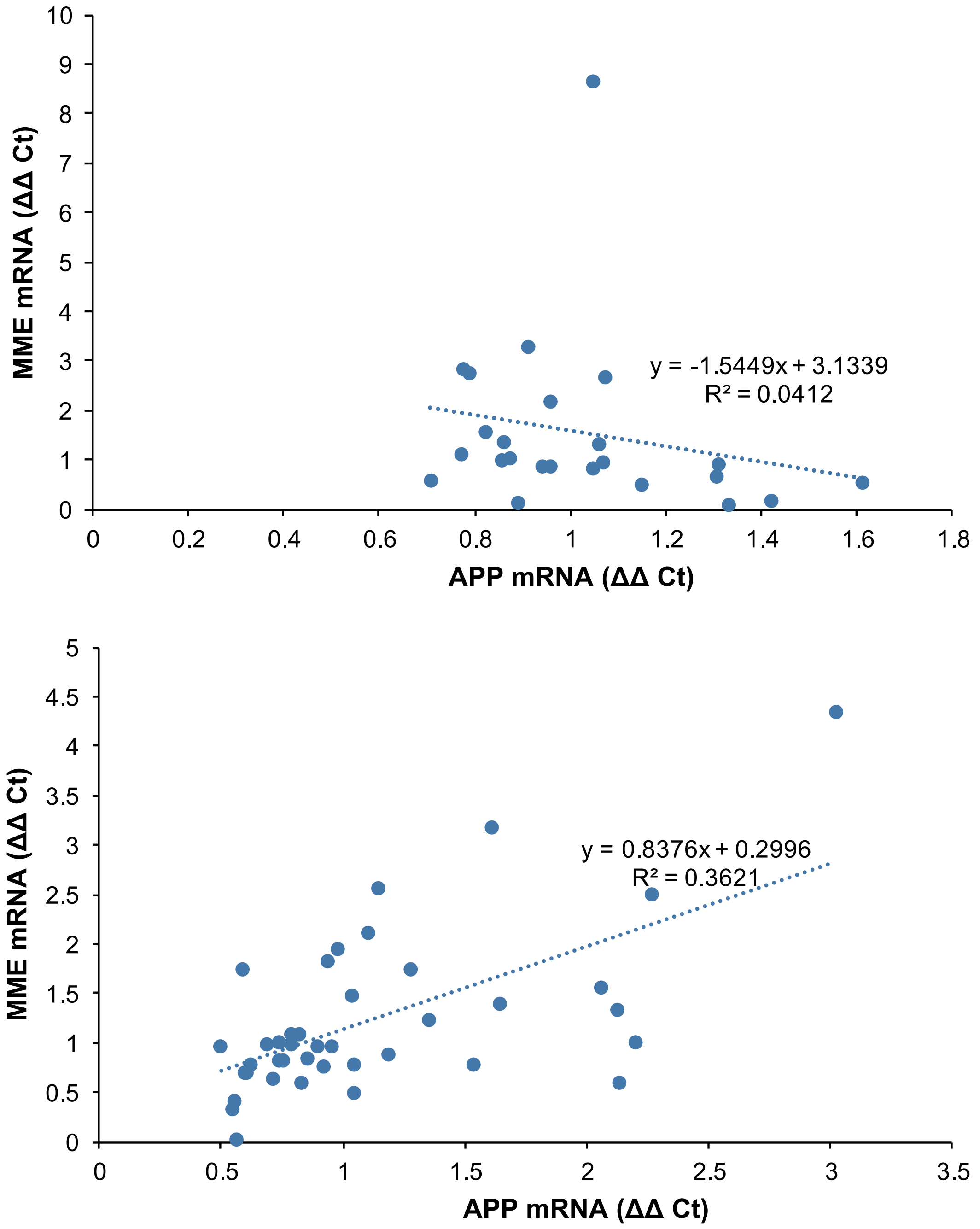
MME mRNA (ΔΔ Ct)
APP mRNA (ΔΔ Ct)
y = -1.5449x + 3.1339
R² = 0.0412
MME mRNA (ΔΔ Ct)
APP mRNA (ΔΔ Ct)
y = 0.8376x + 0.2996
R² = 0.3621

# S3

A. MAPT 3'-UTR reporter clone

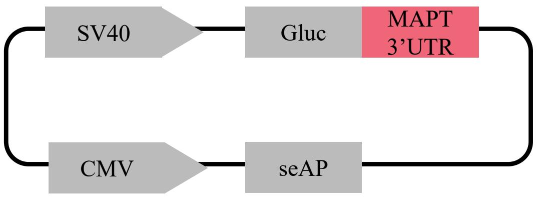


B. MAPT 5'-UTR reporter clone

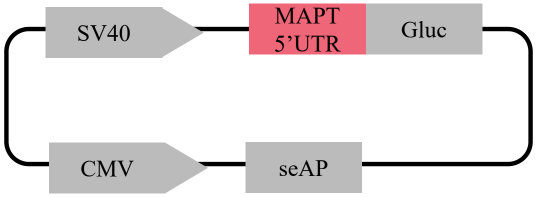


C. MAPT 3'-UTR activities

| ANCOVA | T ratio | P |
|---|---|---|
| plasmid | 237.7 | **<0.0001*** |
| Treatment | 18.11 | **<0.0001*** |
| Plasmid x Treatment | 5.25 | **0.0006*** |
| Plasmid [MAPT 5'UTR] x Treatment [miR-181d] | 0.72 | 0.4729 |

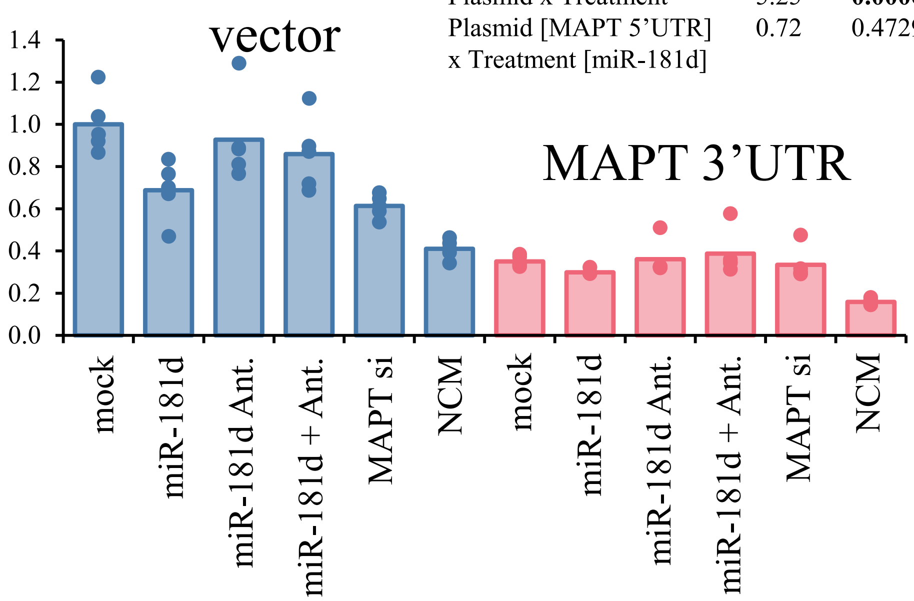


D. MAPT 5'-UTR activities in HeLa

| ANCOVA | T ratio | P |
|---|---|---|
| plasmid | 1.23 | 0.2728 |
| Treatment | 2.64 | **0.0346*** |
| Plasmid x Treatment | 2.56 | **0.0393*** |
| Plasmid [MAPT 5'UTR] x Treatment [miR-181d] | -2.82 | **0.0070*** |

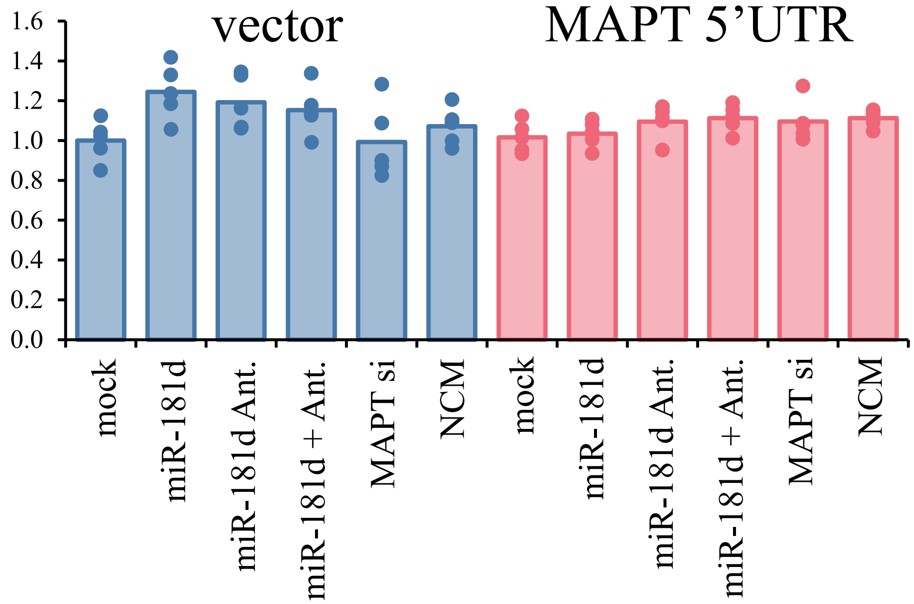


E. MAPT 5'-UTR activities in SK-N-SH

| ANCOVA | T ratio | P |
|---|---|---|
| plasmid | 15.0 | **0.0003*** |
| Treatment | 0.78 | 0.5722 |
| Plasmid x Treatment | 0.94 | 0.4650 |
| Plasmid [MAPT 5'UTR] x Treatment [miR-181d] | -0.70 | 0.4854 |

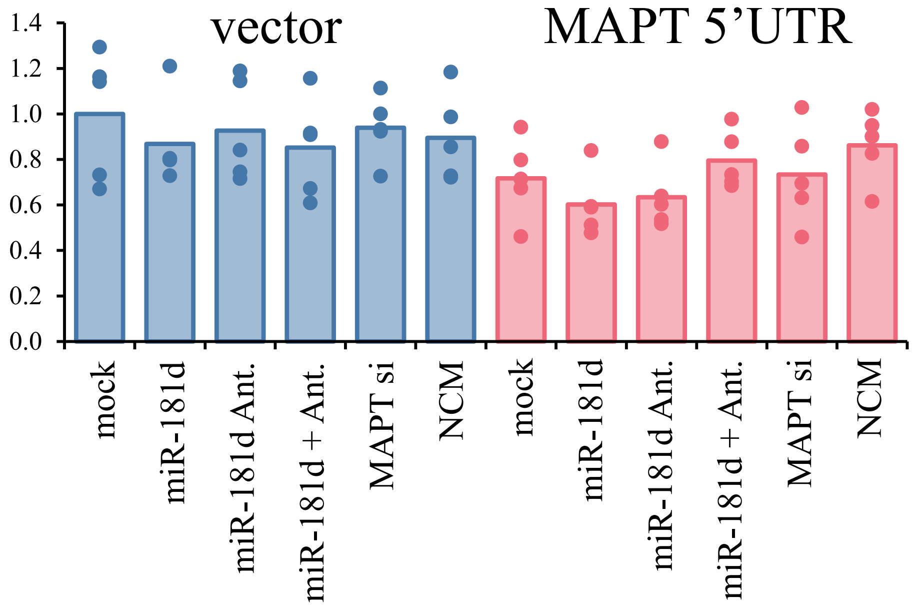

# S6

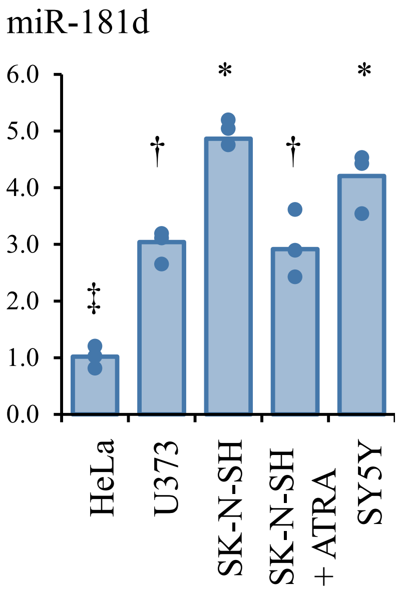
